# Leveraging long-read assemblies and machine learning to enhance short-read transposable element detection and genotyping

**DOI:** 10.1101/2025.02.11.637720

**Authors:** Austin Daigle, Logan S. Whitehouse, Roy Zhao, JJ Emerson, Daniel R. Schrider

## Abstract

Transposable elements (TEs) are parasitic genomic elements that are ubiquitous across the tree of life and play a crucial role in genome evolution. Advances in long-read sequencing have allowed highly accurate TE detection, though at a higher cost than short-read sequencing. Recent studies using long reads have shown that existing short-read TE detection methods perform inadequately when applied to real data. In this study, we use a machine learning approach (called TEforest) to discover and genotype TE insertions and deletions with short-read data by using TEs detected from long-read genome assemblies as training data. Our method first uses a highly sensitive algorithm to discover potential TE insertion or deletion sites in the genome, extracting relevant features from short-read alignments. To discriminate between true and false TE insertions, we train a random forest model with a labeled ground-truth dataset for which we have calculated the same set of short-read features. We conduct a comprehensive benchmark of TEforest and traditional TE detection methods using real data, finding that TEforest identifies more true positives and fewer false positives across datasets with different read lengths and coverages, while also accurately inferring genotypes and the precise breakpoints of insertions.

By learning short-read signatures of TEs previously only discoverable using long reads, our approach bridges the gap between large-scale population genetic studies and the accuracy of long-read assemblies. This work provides a user-friendly tool to study the prevalence and phenotypic effects of TE insertions across the genome.

## INTRODUCTION

Transposable elements (TEs) are genetic sequences capable of replicating themselves throughout the genome and that shape genome structure, gene expression, and evolutionary dynamics (Bourque et al. 2018; Drongitis et al. 2019; Makałowski et al. 2019; Wells and Feschotte 2020). TE insertions can impact phenotypes through the direct disruption of DNA sequences (Finnegan 1992), TE-induced chromosomal rearrangements (Montgomery et al. 1987; Montgomery et al. 1991), and changes in gene expression (Feschotte 2008; Lee 2015). Though most insertions leading to phenotypic changes are thought to be deleterious (Charlesworth and Langley 1989; Lynch 2007), numerous examples of adaptations driven by TE insertions have accumulated (reviewed in Casacuberta and González 2013; Schrader and Schmitz 2019).

To comprehensively quantify the impact of TEs on evolutionary processes, studies usually rely on tools that can detect and genotype TE insertions using genome re-sequencing data. Although recent breakthroughs in long-read sequencing have dramatically improved our ability to pinpoint and characterize TE insertions with unprecedented detail (Kirov et al. 2021; Han et al. 2022; Hoyt et al. 2022; Rech et al. 2022; Mohamed et al. 2023), these technologies remain more costly and less scalable than their short-read counterparts for the time being.

Consequently, most large-scale population genetic studies rely heavily on short-read data, requiring robust computational methods for detecting and genotyping TEs from these more accessible, but inherently limited, resources.

Though numerous TE detection methods have been developed since the advent of whole-genome sequencing, most TE callers that use short-read sequence data (e.g. Illumina) follow a similar general strategy: identifying discordant read pairs, where one read maps to the reference genome and the other to a TE sequence, to identify locations of TE insertions (see Figure 1A).

**Figure 1:**
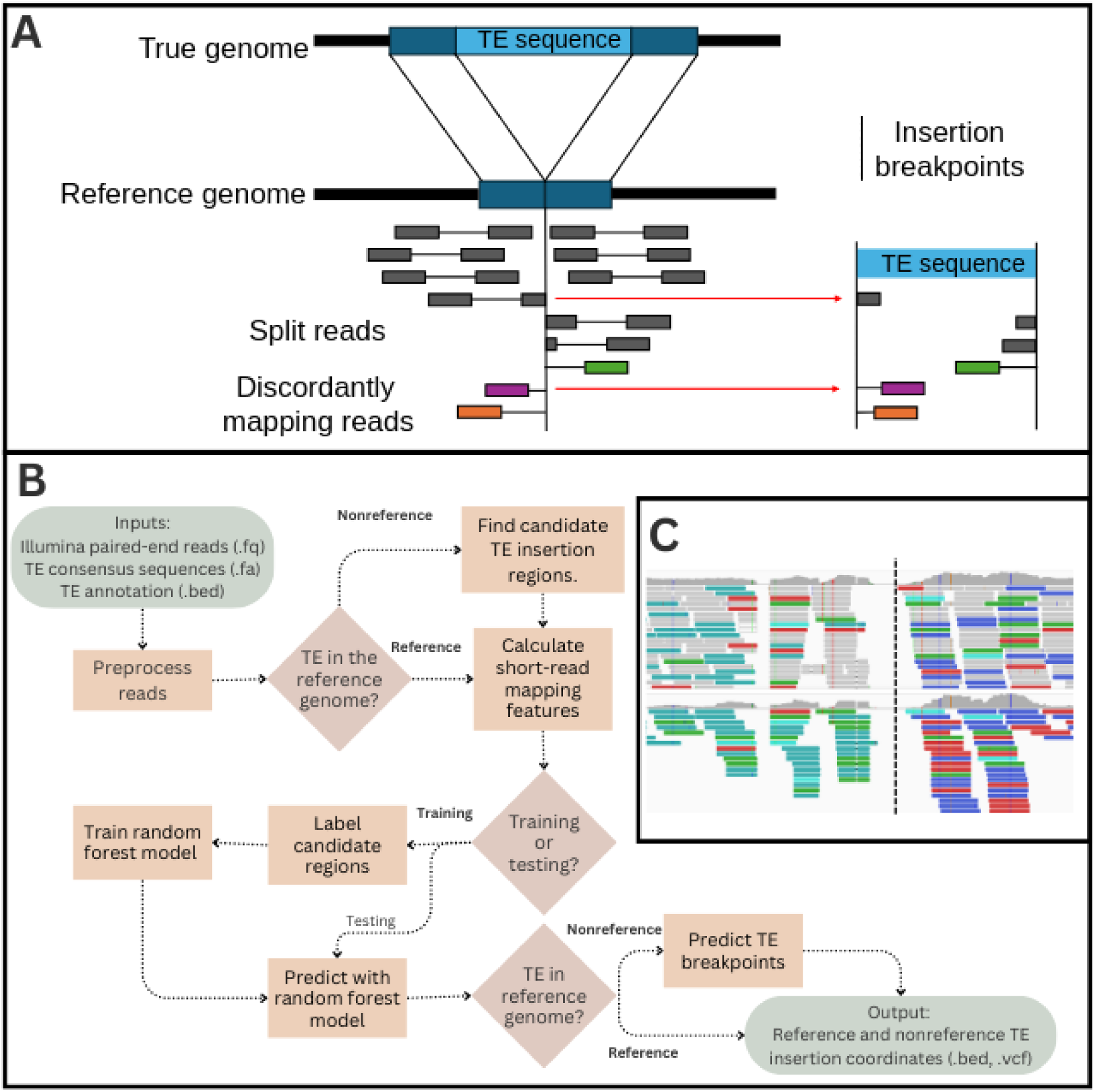
(**A**) Diagram depicting read-mapping information used to detect non-reference TE insertions with paired-end short reads. Short reads are aligned to the reference genome, and reads where one read in the pair maps to the reference genome and the other to a TE sequence (discordantly mapping reads) or reads where one pair is split between the reference and TE sequence (split reads) are quantified. (**B**) Overview of the TEforest pipeline. Input and output files are shown in ovals, important branching points in the pipeline are shown in diamonds, and computational steps of the pipeline are shown in rectangles. (**C**) An example of short-read alignment patterns around a TE insertion, displayed in IGV. Reads mapping to a TE sequence elsewhere in the genome are shown in colors.

Additionally, some methods also use split reads, where part of a single read maps to the reference genome and another portion of that read shares homology with a TE, to precisely identify TE insertion breakpoints. This process is followed by a series of filtering steps to limit the prediction of false positives (reviewed in Makałowski et al. 2019). These approaches can be complicated by the difficulty of designing filters that apply to all read lengths and insert sizes, work for diverse TE family sequences, and do not filter out TE insertions in repetitive regions or nearby other structural variations relative to the reference genome. While these approaches have proven useful, their accuracy often varies substantially according to the properties of the input data, including read length, sequencing coverage, or the degree of repetitiveness in the genome of the species being examined (Vendrell-Mir et al. 2019; Yu et al. 2021; Verneret et al. 2024).

Furthermore, because most previous studies comparing the performance of TE detectors relied on benchmark sets of simulated TE insertions (Nelson et al. 2017; Chen et al. 2023; Verneret et al. 2024) or a small number of genomes assembled with long reads (Rishishwar et al. 2017; Vendrell-Mir et al. 2019; Yu et al. 2021) to perform these assessments, our understanding of the shortfalls and relative efficacy of TE detection methods is incomplete. Simulated data in particular may not adequately represent the complexity and messiness of short-read mapping patterns around true TE insertions. On the other hand, large high-quality datasets of known TE insertions such as the long-read based population-scale *Drosophila melanogaster* assemblies and TE annotations generated by Rech *et al*. (2022), open the possibility for more rigorous testing of TE detection algorithms. Crucially, when both long-and short-read data are available for the same genomes, short-read-based detection methods can also be benchmarked using a set of high-confidence TE insertions. Moreover, the availability of such data allows us to reframe short-read TE detection as a machine learning problem—rather than designing a method to detect what we think the patterns of short-read mapping around a TE insertion *ought* to look like, we can train a machine learning classifier to detect the actual signatures of known TE insertions in real empirical datasets.

Here, we present TEforest, a machine learning method that enhances short-read TE detection and genotyping by learning predictive features directly from high-confidence TE insertions previously detected from long-read assemblies. First, TEforest uses a sensitive initial scanning algorithm to identify a large set of potential TE insertions. TEforest then employs a random forest classifier to simultaneously discriminate between true and false TE candidates and genotype the insertions as heterozygous or homozygous. By training TEforest to examine a rich suite of features—drawn from multiple read mapping signatures and tested across variable read lengths and coverages—we are able to not only achieve higher performance than existing tools, but also provide more reliable genotype predictions, precise breakpoint predictions, and accurate allele frequency estimates. TEforest is freely available at https://github.com/SchriderLab/TEforest.git.

## METHODS

### Algorithm overview

TEforest accepts as input (1) paired-end short-read fastq files, (2) a reference genome in fasta format, (3) a TE consensus library in fasta format, and (4) a BED file detailing reference TE locations (Figure 1B). The algorithm first identifies genomic regions that may contain non-reference TE insertions by finding reads that map to TE consensus sequences and TEs annotated in the reference genome. After a small number of filters are applied to the candidate insertion sites, a comprehensive set of features summarizing read alignments within each candidate region are computed and transformed into feature vectors. These vectors are then classified by a random forest model as either a homozygous TE insertion, a heterozygous TE insertion, or no insertion. These feature vectors can also be used for training a model if the true genotypes are available.

Finally, the algorithm attempts to pinpoint precise breakpoint locations using split-read evidence. For TEs annotated in the reference genome, we use an additional random forest model trained to detect presence/absence using the same feature vectors of the non-reference model.

### Algorithm to detect non-reference insertions

#### Discovery of regions with candidate TE insertions

To discover regions of the genome with potential TE insertions, the fastq reads are preprocessed with *fastp* to trim adaptors and low-quality sequences (Chen 2023) and then mapped to the TE consensus sequences as well as sequences annotated as TEs in the reference genome using the BWA-MEM 2 algorithm, with default settings (Li and Durbin 2009; Vasimuddin et al. 2019).

Reads that map to TE sequences are then mapped back to the reference genome using the same software/settings. Nested TE sequences, where an annotation of one TE overlaps with another TE, are omitted to prevent errors where sequences aligning to one TE are misattributed to another nested TE. For each TE family, the locations of read pairs where one of the two reads maps to the TE sequence (whether it be the canonical sequence or an annotated copy of that TE family found in the reference genome) are used to identify candidate regions for TE insertions; we refer to these pairs as “TE-mapping read pairs”. Initially, the candidate region consists of any contiguous stretch of sites with coverage by a TE mapping read. To avoid confounding reads mapped to reference TEs with non-reference insertions, candidate regions overlapping with a reference TE of the same family as well as those contained entirely within a reference TE of a different family are filtered out.

Because we observed that many false positive regions are very short in length, regions <154 bp are also filtered out to improve computational efficiency. This length is slightly larger than the largest read length used in our dataset and was chosen because many of these false positive regions are the length of a single read. Shortening this filtering length did not improve performance for shorter read lengths (not shown). To ensure that our candidate region fully encompassed all TE-mapping read pairs around a putative TE insertion, all remaining regions are expanded a further 200 bp in either direction. Because many TE insertions contain gaps in coverage of TE-mapping read pairs (Figure 1C), we next merge any overlapping candidate regions for the same TE family. The implementation of this algorithm benefitted from open source bioinformatics packages for working with short-read alignments and genomic coordinate ranges including SAMtools (Danecek et al. 2021), SeqKit2 (Shen et al. 2024), and GenomicRanges (Lawrence et al. 2013).

#### Calculation of feature vectors

Here we expand on concepts described by Hill and Unckless (2019), who proposed the use of a feature vector as input for machine learning models to detect structural variants. This feature vector representation summarizes many aspects of the alignment of short reads to the genome calculated on a per-base level (all features are defined in Table 1). Features are calculated based on the alignment of all reads to the reference genome as well as to a TE family-specific BAM file, which contains the alignments of TE-mapping read pairs discovered during the candidate region discovery step. For each site in the candidate region a per-base pair sum is calculated for all features listed in Table 1 by extracting information from the CIGAR string, bitwise flag, coverage, and other alignment information reported by the BAM file containing the alignment of all reads to the reference genome. For each site, the value of each of these features is divided by the by the total number of reads mapping to that site. A concise vector summary of these features for a candidate region is created using the mean, standard deviation, median, and interquartile range (IQR) for each statistic across all sites in the candidate region, resulting in 21×2×4=168 feature summaries in total (21 features measured for both TE-and reference-genome mapping read pairs, and each summarized by four values) in the final vector.

**Table 1:** Description of features used to describe the alignment of short reads to the reference genome. Each feature is first calculated for each basepair of each read and summed for each basepair before summary statistics (mean, median, sd, and IQR) are calculated for the feature across the region. Features are calculated using the full BAM alignment as well as a BAM file containing only reads aligning to the TE of interest.

| Feature | Description (T=1, F=0) |
| --- | --- |
| Cigar1 | Base matches reference. |
| Cigar2 | Deletion has occurred in the read. |
| Cigar3 | Insertion in one or more bases in the read, preceding this base. |
| Cigar4 | Soft clip in the read at this position. |
| Cigar5 | Hard clip in the read at this position. |
| Paired | Read has a pair. |
| Proper Pair | Both reads in the pair are aligned with correct orientation and distance. |
| Is Read1 Unmapped | First read in the pair is unmapped. |
| Is Read2 Unmapped | Second read in the pair is unmapped. |
| Is Read1 Rev Comp | First read in the pair is reverse complemented. |
| Is Read2 Rev Comp | Second read in the pair is reverse complemented. |
| Is First Read | Is the first read in the pair. |
| Is Second Read | Is the second read in the pair. |
| Split | Read is part of a split alignment. |
| Long Insert | Template length of this read is in the top 1% of template lengths in this region. |
| Short Insert | Template length of this read is in the bottom 2% of template lengths in this region. |
| Parallel Read | Both read and its mate align in the same direction. |
| Everted Read | Both read and its mate align in opposite directions. |
| Discordant Read | Mate of the read is not mapped to the same chromosome. |
| Template Length | Distance between the two ends in the read pair on the reference genome. |
| Quality | Quality score of the alignment at this position. |

#### Random forest model

When the locations of true TE insertions are known, labeled feature vectors can be used for training a machine learning model. We used a random forest classifier, implemented in scikit-learn (Pedregosa et al. 2011), and trained with 500 estimators (i.e. decision trees), using the Gini impurity criterion to measure split quality. To address potential class imbalance, we set class_weight=“balanced”, which adjusts weights inversely proportional to class frequencies. The number of features to consider when looking for the best split was set to the square root of the total number of features. All other hyperparameters were set to their default values in scikit-learn. The possible output classes for the classifier are homozygous insertion, heterozygous insertion, or no insertion.

#### Breakpoint detection

After the random forest model described above is used for TE detection, all candidate regions classified as homozygous and heterozygous insertions are further processed to identify precise breakpoints of TE insertions. TE-mapping read pairs are checked for split reads, and if none are identified, then all read pairs mapping to the candidate region are checked for split reads. For each split read, we locate the site in the candidate region where the 3’-most site in the read maps to before the split, calling this a split-read-end. We find the top two sites with the largest number of split-read-ends. We use these two sites as the final TE breakpoints, as most TEs make staggered double strand breaks when inserting into the genome, creating a target site duplication at the insertion site (Craig 2007; Linheiro and Bergman 2012). If only one site with split reads is identified, that site is used as the insertion site. If no split reads are identified, then the center of the candidate region is used as the breakpoint.

### Algorithm to infer the presence/absence of TE insertions annotated in the reference genome

When examining short-read data from a given individual, TEforest classifies TEs annotated in the reference genome as present or absent in a similar manner to non-reference insertions, though without a genotyping step. For this task, no detection of putative candidate regions or breakpoints is needed. User-provided coordinates of reference TEs are expanded 500 basepairs in either direction to capture information about reads surrounding the TE breakpoints. Feature vectors are calculated as previously described, and a random forest classifier is trained and used to label a reference TE as present or absent using the same hyperparameters used for classifying non-reference TEs.

### Testing TEforest on high-quality *Drosophila melanogaster* data

#### Creation of synthetic heterozygous genomes for training/testing TEforest’s genotyping of non-reference TEs

A subset of the *Drosophila melanogaster* genomes annotated with the manually curated TE library created by Rech *et al*. (2022) were used for training and testing the TEforest model. As this annotation only contains homozygous insertions found in the long-read assembly, we selected six genomes with low heterozygosity (as reported in Supplemental Table 2 of Rech *et al*. (2022) to minimize the risk of our model identifying false positive insertions that are present in only a subset of flies from the inbred line used for DNA extraction and sequencing.

To include known heterozygous insertions in model training, we created synthetic heterozygous genomes by combining reads from two genomes sequenced with the same read length. To do this, reads from both genomes were first preprocessed by *fastp* then mapped to the reference genome. A final heterozygous fastq file was created such that half the reads from contigs 2R and 3R were taken from each genome, while all other chromosome arms were taken from one of the genomes in the pair, with reads in each chromosome arm downsampled to the same mean coverage. Thus, most differences between the two lines on chr2R and chr3R would be heterozygous, while all variants on the remaining chromosome arms would be homozygous. To train on a diverse set of read lengths, we created synthetic heterozygous genomes from genomes sequenced with 54, 125, and 151 bp reads (one synthetic genome per read length).

#### Training and validation dataset for non-reference insertions

TEforest was trained using non-reference insertions from contigs 3L, 3R, and X and tested on contigs 2L and 2R to ensure that no insertions used in model training were used in model testing. Additionally, all candidate regions found inside the heterochromatic regions were filtered out as these regions were not annotated by Rech *et al*. (2022). A full description of the numbers of labeled candidate regions used for training and testing can be found in Table 2. Separate models were trained and tested at target coverages of 5X, 10X, 20X, 30X, 40X, and 50X.

**Table 2:** Numbers of homozygous, heterozygous, and false positive candidate regions used as input for training and testing the non-reference insertion model (50X coverage).

| Training/testing | Read length (bp) | Insert size (bp) | # homozygotes | # heterozygotes | # candidate regions without true positives |
| --- | --- | --- | --- | --- | --- |
| Training | 54 | 287 | 204 | 252 | 778 |
| Training | 125 | 208 | 200 | 272 | 25863 |
| Training | 151 | 436 | 202 | 312 | 4080 |
| Testing | 54 | 287 | 129 | 219 | 255 |
| Testing | 125 | 208 | 110 | 192 | 10390 |
| Testing | 151 | 436 | 108 | 218 | 1586 |

#### Training and validation data for reference insertions

A model to detect reference TE presence and absence was trained using highly homozygous genomes only because reference insertions tend to be homozygous present or absent. Reads were downsampled to the same target coverages and training and testing was conducted on the same contigs as the non-reference model. The numbers of presence and absence calls found in training and testing can be found in Table 3.

**Table 3:** Numbers of candidate regions with a TE present or absent used as input for training and testing the reference insertion model (50X coverage).

| Training/testing | Read length (bp) | Insert size (bp) | # present | # absent |
| --- | --- | --- | --- | --- |
| Training | 54 | 287 | 1668 | 1213 |
| Training | 125 | 208 | 1681 | 1219 |
| Training | 151 | 436 | 1554 | 1328 |
| Testing | 54 | 287 | 374 | 320 |
| Testing | 125 | 208 | 389 | 319 |
| Testing | 151 | 436 | 333 | 362 |

### Benchmarking other TE detectors

To compare TEforest to other TE detectors, we utilized McClintock, a meta-pipeline that implements popular TE detection methods in a controlled workflow (Nelson et al. 2017; Chen et al. 2023). We ran McClintock with default settings, using PoPoolationTE (Kofler et al. 2012), PoPoolationTE2 (Kofler et al. 2016), RetroSeq (Keane et al. 2013), TEFLoN (Adrion et al. 2017), TEMP (Zhuang et al. 2014), and TEMP2 (Yu et al. 2021). These methods were chosen based on their ability to produce more than zero calls for all read lengths tested and finish running within five days using 8 CPU cores.

Methods were benchmarked for accuracy of TE detection, genotyping, and breakpoint localization. To assess the overall ability to detect TEs, calls made by a TE caller within 500 bp of a TE of the same family in the truth dataset were counted as true positives, with the rest being counted as false positives. Overlapping TEs of the same type in the truth dataset were condensed to avoid the double-counting of nested TEs; this was not required for nested TEs of different types. The truth dataset provided by Rech *et al*. (2022) does not provide perfectly precise breakpoints because it does not report the target site duplications (TSDs) produced by the staggered cuts of transposase upon TE integration (Craig 2007; Linheiro and Bergman 2012); rather, it provides an approximate breakpoint near the two breakpoints of the TSD. For this reason, the center of two breakpoints predicted by each TE caller was used to calculate distances between the truth dataset breakpoint and the prediction of the TE caller. If one breakpoint is predicted, we simply found the distance between this breakpoint and the truth dataset breakpoint. To assess genotyping accuracy, we calculated the *F*_1_ scores separately for homozygous and heterozygous genotypes and reported the macro-averaged *F*_1_ score as the overall measure of performance. For example, for the homozygous *F*_1_ score, the true positives are correctly genotyped homozygotes, false positives are heterozygotes classified as homozygotes, and false negatives are homozygotes classified as heterozygotes, and vice-versa for the heterozygous *F*_1_ score. We additionally performed a separate benchmarking of breakpoint accuracy and genotyping performance for the intersection of calls made by TEforest, TEMP2, and RetroSeq at a given coverage.

### Benchmarking allele frequency accuracy

To assess the ability of TE callers to accurately predict the allele frequencies of non-reference TEs in a population of individuals, we benchmarked the performance of all previously used TE callers in 13 genomes from the *Drosophila* Synthetic Population Resource, sequenced with 54 bp reads, that were included in our truth dataset. The median coverage of these genomes was 46X, so the TEforest model trained with 50X coverage was used for this task. Only calls within the euchromatic regions of contigs 2L, 2R, 3L, 3R, and X were used for benchmarking. As these genomes are highly homozygous, each true positive call (a prediction of a TE insertion of the same family within 500 bp of an annotated insertion site) was counted as a frequency of one, with a max frequency of 13. For each insertion in the population, the true allele frequency was subtracted from the predicted allele frequency.

## RESULTS

### TEforest outperforms other TE detection methods

We developed TEforest, a machine-learning tool that detects the positions of transposon insertions in a reference genome using short reads (Figure 1 and Methods). This is accomplished by identifying read pairs where one read maps to an annotated set of TE sequences and the other to the reference genome (TE-mapping read pairs), extracting a comprehensive set of features describing read-mapping patterns in those genomic regions containing TE-mapping read pairs, then using a random forest algorithm to predict if the region contains a true TE insertion. By comprehensively incorporating all available read mapping information from both true TE insertions and non-insertion sites, our approach attempts to capture a comprehensive spectrum of read-mapping patterns, thereby enabling the model to more effectively distinguish genuine insertions from false positives. We trained and tested TEforest using the annotated TE insertions in six *D. melanogaster* genomes sequenced with both long and short reads by Chakraborty *et al*. (Chakraborty et al. 2019) and Rech *et al*. (2022). Because these genomes were sequenced from inbred lines and we wished to train TEforest to be able to detect and genotype both homozygous and heterozygous TE insertions, we created synthetic heterozygous non-reference TE insertions by combining the reads of highly homozygous genomes for some chromosome arms, while others remained homozygous. These data were partitioned into training and testing sets as described in the Methods and in Table 2. In addition to benchmarking TEforest on this hold-out test set, we benchmarked six other TE detectors available in the McClintock meta-pipeline.

Overall, TEforest demonstrated better accuracy than competing methods across our three tested read lengths and varying coverages (Figure 2A-C; SFigure 1&2A-C). TEforest’s superior performance is largely due to an increase in recall, as most TE detectors including TEforest called very few false positives. In general, the *F*_1_ score (the harmonic mean of recall and precision) increased with coverage for all callers due to an increased number of informative reads. Interestingly, TEforest tended to approach its highest *F*_1_ score (∼0.84) at low coverages (10-30X) before leveling off, while other callers continued to show slight increases up to 50X coverage. However, no method achieved recall above 75% for any coverage, indicating that a subset of TEs was undetectable with any amount of short-read data (Figure 2D; SFigure 1&2D). In the case of TEforest, the majority of these false negatives (69%) were lost in the candidate region detection stage instead of being mislabeled by the random forest model, which could be due to a lack of supporting reads or proximity to an annotated reference TE resulting in the region being filtered out (Methods).

**Figure 2:**
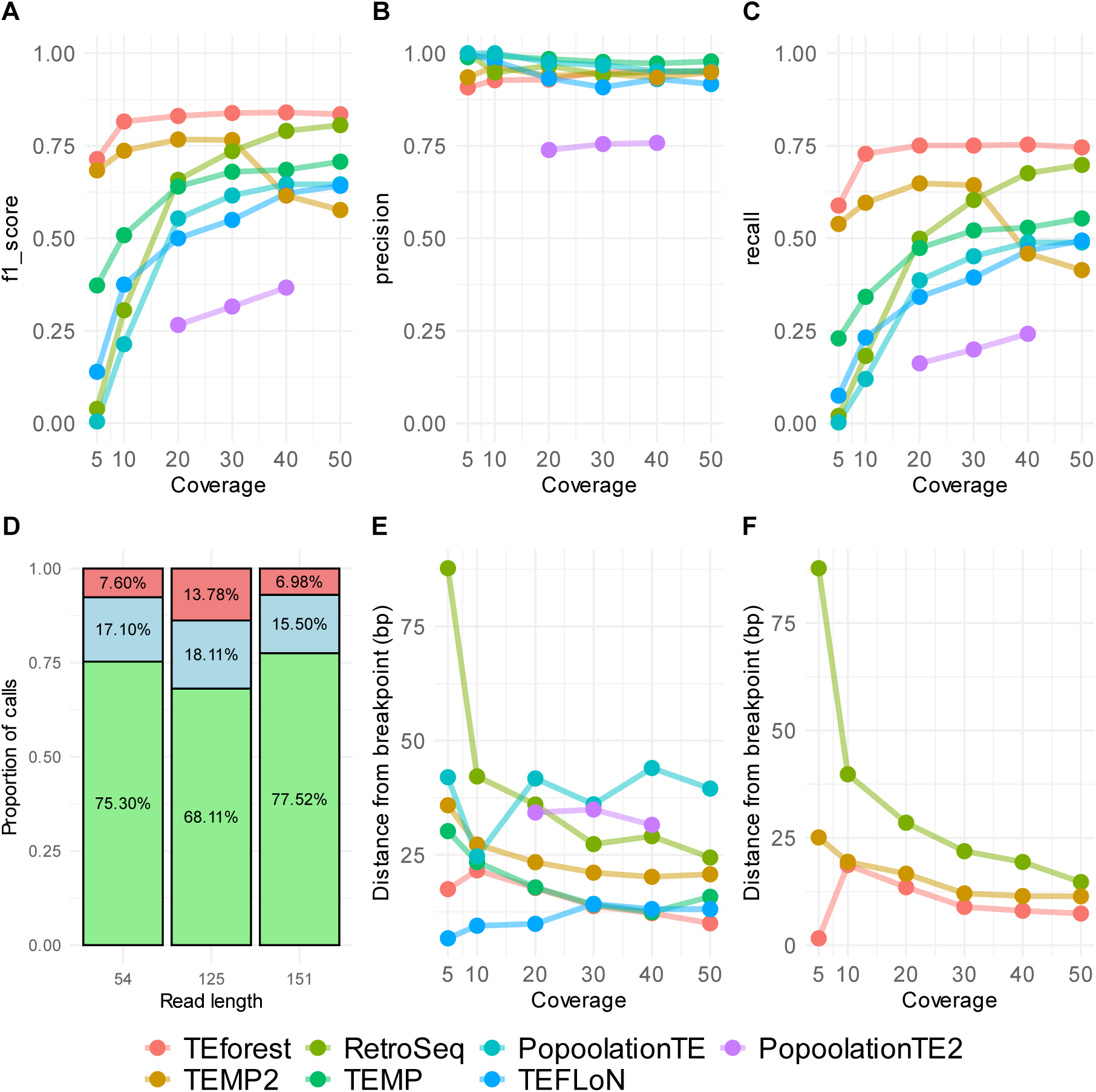
The performance of TEforest compared to other short-read TE callers at detecting non-reference TE insertions annotated by long read assemblies of *D. melanogaster* strains. Short reads used for detection of TEs were 151 bp with ∼436 bp insert sizes. True positives were defined as predicting a TE insertion of the correct TE family with any prevalence within 500 bp of the true insertion site. Performance was quantified with (**A**) *F*_1_ scores, (**B**) precision, and (**C**) recall. The proportions of TEs that were successfully detected by TEforest, lost in candidate region detection stage, or misclassified by the random forest model for each read length are shown in Panel **D**. The mean breakpoint accuracy of TE callers for all (**E**) true positive calls or (**F**) true positive calls shared by TEforest, RetroSeq and TEMP2 was quantified by finding the distance between the center of true positive breakpoint ranges predicted by the TE callers and the breakpoint in the truth dataset.

The other TE callers tested exhibited more variable performance than TEforest across different read lengths and coverages. When tested with 151 bp reads, TEMP2 had the next-best *F*_1_ score compared to TEforest from 5X-30X coverage (e.g. 0.68 at 5X coverage and 0.77 at 30X coverage, vs. 0.71 at 5X coverage and 0.84 at 30X coverage for TEforest), before seeing a rapid decline in recall at 40X (leading to a *F*_1_ score of 0.62 for TEMP2 vs. 0.84 for TEforest) and 50X coverage (*F*_1_ of 0.58 for TEMP2 vs. 0.84 for TEforest). RetroSeq continued to steadily increase in recall, eventually approaching but not exceeding the performance of TEforest at 50X coverage (*F*_1_=0.81).

When tested with 125 bp reads, TEMP2 was consistently the next-best caller behind TEforest (TEforest’s *F*_1_=0.80 at 50X), though it still exhibited a decline in performance when increasing coverage to 50X (*F*_1_=0.76 at 40X and 0.70 at 50X; SFigure 1), although this was not as severe as in the 151 bp data set. In contrast to the 151 bp dataset, RetroSeq’s performance lagged behind the other TE callers until 50X coverage (*F*_1_=0.49 at 40X and 0.59 at 50X), and did not approach the *F*_1_ score of TEforest. Interestingly, the median insert size for this dataset was 208, meaning that the paired end reads typically had no gap between them when aligned to the genome. As RetroSeq only utilizes information from discordant paired-end reads rather than information about split reads for TE detection, it may struggle to detect TEs in this context. In contrast to the 151 and 125 bp datasets, when tested on the 54 bp dataset, TEforest exhibited lower recall and *F*_1_ scores at 5X and 10X coverage (*F*_1_=0.58 at 5X) compared to TEMP and TEMP2, which maintained their high *F*_1_ scores at even 5X coverage (*F*_1_=0.80 and 0.83, respectively; SFigure 2). From 20X-50X coverage, *F*_1_ scores were approximately equal among these callers (∼0.85). We also note that in the 54 bp dataset TEMP2 did not exhibit any of the decline in performance at higher coverages that it did in the 151 and 125 bp data sets.

### TEforest non-reference calls are close to the true insertion breakpoints

We additionally approximated the breakpoint accuracy of TEforest and other TE detectors by measuring the mean absolute difference between the true and predicted insertion sites for each method. For the 151 bp dataset, TEforest’s mean breakpoint estimation error was <25 bp for all coverages, gradually improving to ∼10 bp at 50X coverage (Figure 2E). TEforest’s predictions were closer on average to the true breakpoint than all callers except for TEFLoN at lower coverages (<30X), and comparable to TEFLoN and TEMP at higher coverages. The full distribution of breakpoint distances from the truth shows that TEforest, TEMP2, and TEFLoN predict ∼50% of calls 0 or 1 bp away from the annotated breakpoint, while an excess of calls >=25 bp away from the annotated breakpoint causes TEMP2 to be 2X further than TEforest from the true breakpoint on average (SFigure 3). RetroSeq had an excess of predictions at intermediate (∼10 bp) and long (>25 bp) distances, which is not unexpected because it does not use split reads to narrow down its breakpoint locations. Callers also improved as the coverage increased for the 125 bp dataset, though nearly all callers performed similarly to each other in this context (SFigure 1E). TEforest produced more distant calls relative to other algorithms in the 54 bp context, likely due to its reliance on split short reads whose paired read maps to a TE, which are less common for shorter reads (SFigure 2E).

As coverage decreased, each method’s recall decreased (although this trend was less severe for TEforest). We hypothesized this may lead to inaccurate comparisons of callers because more difficult to detect TEs may also be more difficult to accurately detect breakpoints for. We compared the intersection of TEs recovered by TEforest, TEMP2, and RetroSeq, and found that TEforest more accurately predicted breakpoint locations than TEMP2 and RetroSeq with 151 and 125 bp reads, but for 50 bp reads was ∼2X further from the true breakpoint relative to TEMP2 and RetroSeq at 30X coverage and above (Figure 2F; SFigure 1&2F). Overall, TEforest can be relied upon to produce calls reasonably close to the breakpoint of a TE insertion.

### TEforest accurately genotypes non-reference TE insertions

We also assessed the ability of TEforest and other callers to accurately genotype known TE insertions. Other callers estimate the fraction of genomes in the sample from which DNA was collected that contain an insertion, yielding a prediction that can range anywhere from 0 to 1. While some tools refer to this as the TE insertion’s “frequency”, to avoid conflation with population frequency we refer to this as the insertion’s “prevalence”. In contrast, since we designed TEforest to classify insertions as heterozygotes or homozygotes, it makes discrete predictions of 0 (homozygous absent) 0.5 (heterozygous) or 1 (homozygous present). Thus, to allow for a comparison of genotyping accuracy among methods, we transformed the predictions made by the other methods into distinct genotypes by labeling estimates below 0.25 as homozygous absent, between 0.25 and 0.75 as heterozygotes, and above 0.75 were called homozygous present.

In the dataset with 151 bp reads at 30X coverage, the predicted prevalence of true positive predictions were located near the true value for TEforest (predicted mean prevalence of 0.98 for homozygous insertions and 0.53 for heterozygous insertions) and RetroSeq (homozygous mean prevalence=0.94, heterozygous mean=0.50), while TEMP2 systematically underestimated the prevalence (homozygous mean=0.46, heterozygous mean=0.18), despite having a high *F*_1_ score of 0.77 for detecting insertions at this coverage (Figure 3A-F). The average prevalence predicted by TEforest for homozygous true positive insertions (that is, the average of its predicted genotypes) remained close to one across all coverages, while it tended to misclassify heterozygotes as homozygotes at lower coverages and make more accurate predictions at higher coverages (Figure 3I). Other callers such as RetroSeq, TEFLoN, and PopoolationTE also had reasonable average prevalences, while TEMP and TEMP2 underpredicted the true prevalence of insertions. We also calculated a genotyping *F*_1_ score, where we compute *F*_1_ scores separately for homozygous and heterozygous genotypes and reported the macro-averaged *F*_1_ score as the overall measure of performance. TEforest displayed a competitive genotyping *F*_1_ score relative to other across callers across coverages when calculated for all of its calls (Figure 3G) or when comparing to the intersection of TE insertions detected by TEforest, RetroSeq, and TEMP2 (Figure 3H). Despite recovering fewer known TEs, PopoolationTE and TEFLoN performed well at the genotyping task for those TEs they were able to detect (Figure 3G).

**Figure 3:**
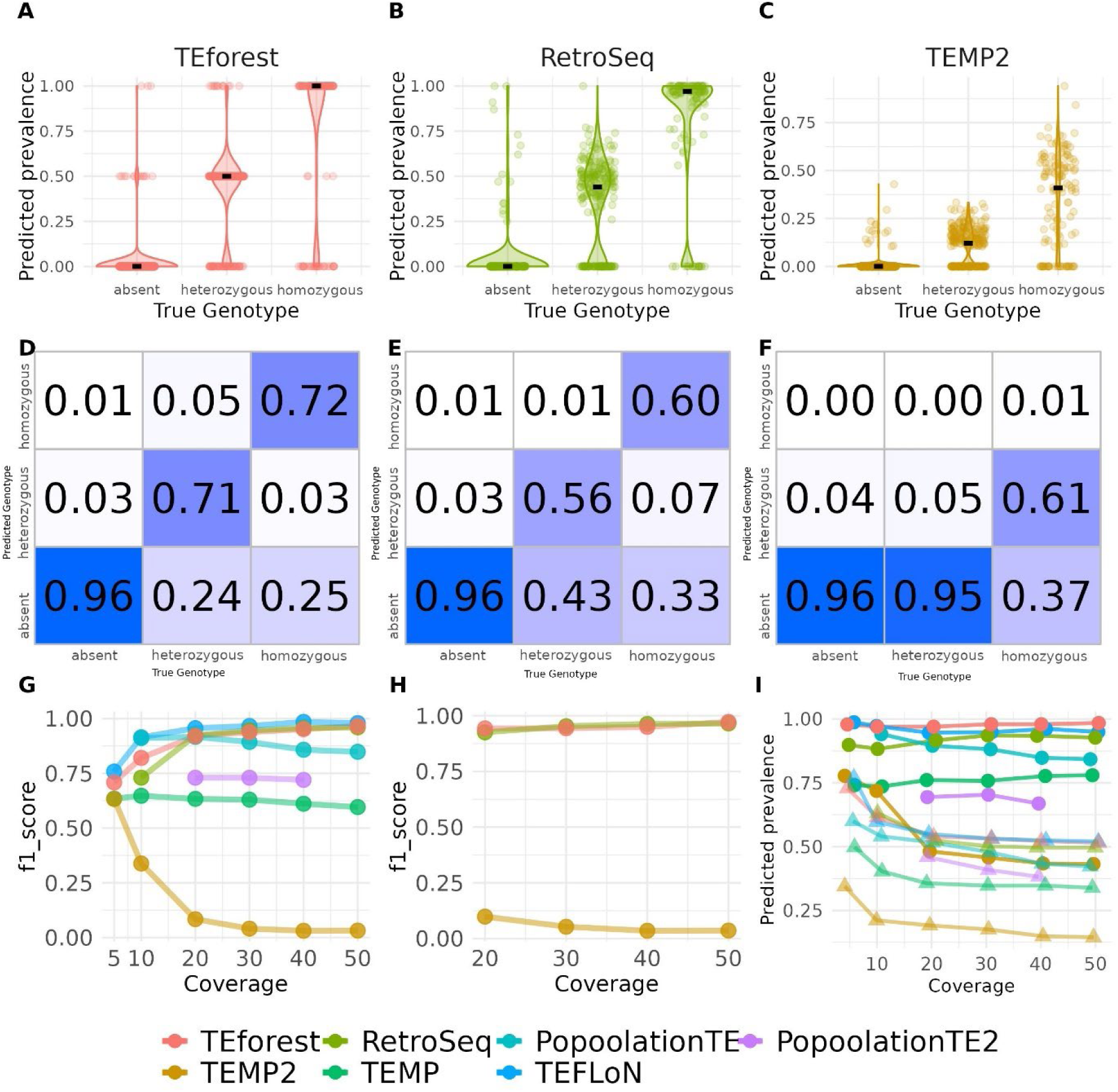
The performance of TEforest compared to other short-read TE callers at genotyping non-reference TE insertions. Short reads used for detection of TEs were 151 bp with ∼436 bp insert sizes. Frequency predictions for insertions that were homozygous, heterozygous, or absent were calculated for (**A**) TEforest, (**B**) RetroSeq, and (**C**) TEMP2, where true positive absences refer to candidate regions identified by TEforest with no TE insertion. The median prevalence is shown as a black line. (**D**-**F**) The accuracy of these frequency predictions is quantified in confusion matrices, where heterozygous predictions are defined as frequency predictions between 0.25 and 0.75 and homozygous predictions are above 0.75. The genotyping *F*_1_ score for (**G**) all true positive calls or (**H**) true positive calls shared by TEforest, RetroSeq and TEMP2 was quantified using true positive predictions of each caller. The mean for the predicted prevalence of true positive homozygous (circles) and heterozygous (triangles) insertions are shown in panel **I**, calculated using only the prevalences of true positive predictions of each caller (false positives are excluded).

For 125 bp reads with short insert sizes, patterns of genotyping were largely the same as for the 151 bp dataset, with TEFLoN, PopoolationTE, and TEforest performing well (SFigure 4). For example, at 30X coverage TEforest’s mean prevalence was 0.96 for homozygotes and 0.55 for heterozygotes. TEMP and TEMP2 also underpredicted the prevalence of insertions at this read length (e.g. for TEMP2 at 30X coverage, homozygous mean prevalence=0.50, heterozygous mean=0.18). RetroSeq, which also performed poorly at detecting known TE insertions, overpredicted the prevalence of heterozygous insertions for this dataset across all coverages (at 30X coverage, homozygous mean=1.00, heterozygous mean=0.77; SFigure 4I). Genotyping performance for all callers was generally higher for the 54 bp reads than other read lengths (SFigure 5), which is to be expected as a more reads are sequenced per genome to reach a given coverage, and the space between the two reads on a sequenced fragment is larger. TEMP and TEMP2 still underpredicted the prevalence of TE insertions, though this bias was less severe than for the longer read lengths (SFigure 5C&I). Specifically, at 30X coverage, the mean prevalence for homozygotes was 0.99, 0.98, and 0.94 for TEforest, RetroSeq, and TEMP2, respectively while the mean prevalence for heterozygotes was 0.52, 0.54, 0.39.

**Figure 4:**
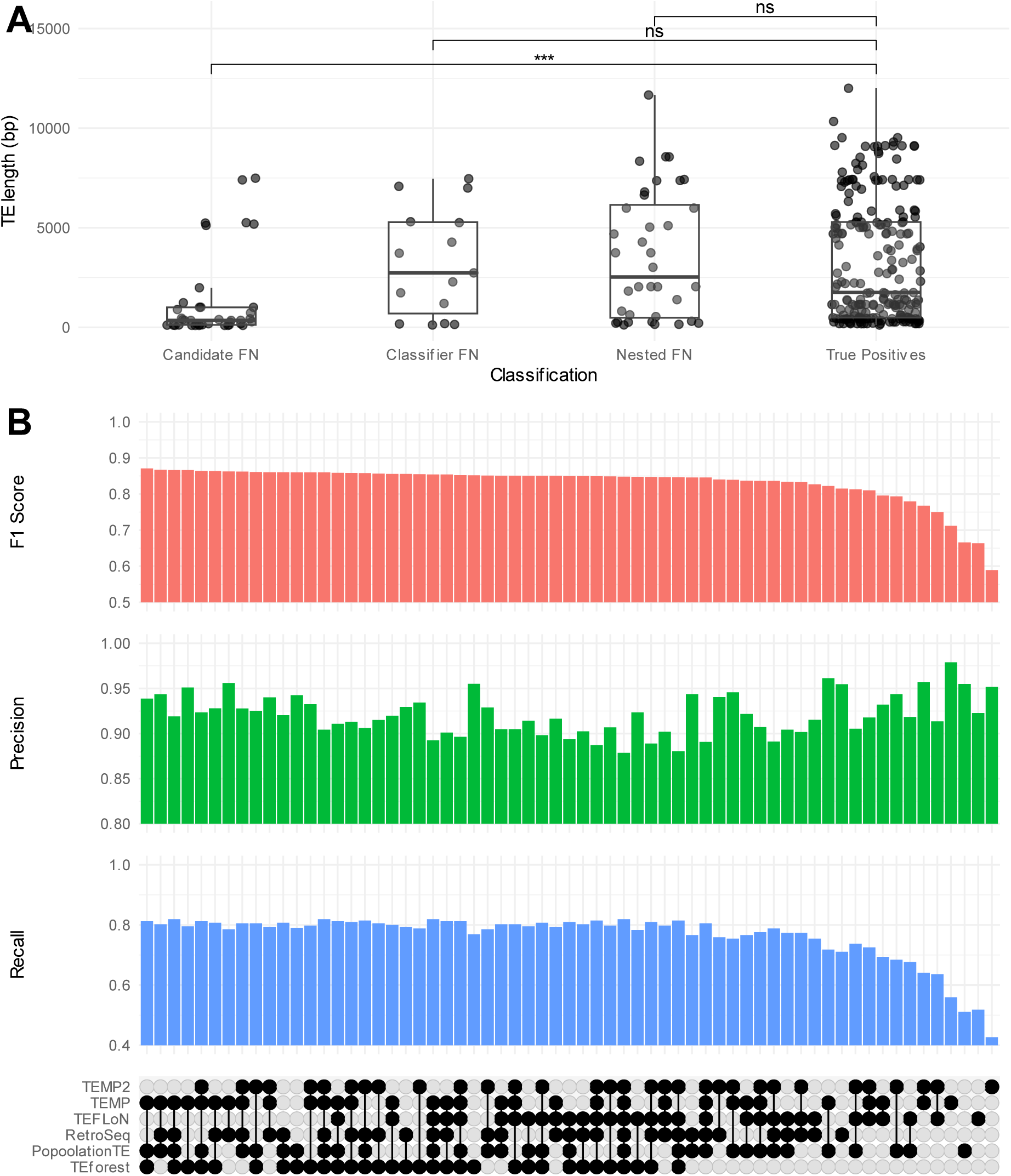
(A) Lengths of non-reference TEs insertions that were in the genome with 151 bp with ∼436 bp insert sizes, divided into classes depending on whether they were unnested and not detected in the candidate region identification stage (Candidate FN), unnested and mislabeled as absences by the random forest classifier (Classifier FN), nested and lost in either the candidate regions or classifier step (Nested FN), or successfully detected by TEforest (True Positives). Overhead bars represent the results of pairwise Wilcox rank-sum tests. **(B)** Upset plot representing the results of combining different TE callers on the *F*_1_ Score, Precision, and Recall for detecting non-reference TEs.

### TEforest false negatives include truncated and nested TEs

Despite its high recall relative to other methods, TEforest still failed to detect a considerable fraction of true positive non-reference insertions (22.5% of the true positive calls in genome with 151 bp reads at 50X coverage). We hypothesized that insertions of TEs that were shorter or more fragmented relative to the consensus TE sequence, nested inside of other TE insertions, and/or located near other structural variants would be more difficult to detect than full-length TE insertions in non-repetitive regions. Focusing on TEforest false negatives for the genome sequenced with 151 bp reads at 50X coverage, we found that insertions that were not detected in the candidate region detection stage of the algorithm were significantly shorter in total length (Figure 4A) compared to their consensus TE sequence (SFigure 6). Lengths of TE insertions were taken from Supplementary Table 10 from Rech *et al*. (2022). Additionally, we found that ∼20% of these false negatives consisted of TEs from the families labeled as INE-1 and Gypsy-2-Dsim in Rech *et al*. (2022), and that most of these insertions from these families were lost in the candidate region stage (STable 1). INE-1 is a TE family that has been thought to be inactive for millions of years (Kapitonov and Jurka 2003; Singh and Petrov 2004), and thus most copies are expected to be old and fragmented, so low performance on this family is not surprising. In addition, TEs nested with copies from different TE families were much more difficult to detect, with 28/37 TEs nested with different TE families not identified by TEforest, with 20 of these lost in the candidate region detection stage (STable 2). TEs nested with copies of the same TE family were easier to detect (though not as easy as unnested TEs), with ∼50% undetected, and all lost in the candidate region detection stage.

### Combining TE insertion detectors improves performance

To determine whether TEforest false negatives were identified by other methods and whether its improved performance stemmed from recovering insertions missed by all other methods, we evaluated the combined performance of various TE detectors in the genome with 151 bp reads at 50X coverage. We found that while TEforest was able to identify twice as many unique true positive insertions as the caller with the next highest number of unique true positives (RetroSeq), ∼96% of true positive insertions found by any caller, including TEforest, were identified by at least one additional caller (SFigure 7A). Notably, of those TE insertions identified by at least one caller, only 6% were not recovered by TEforest.

While all callers produced false positives that were not found by any other callers, 23% of false positives were shared with at least one other caller (SFigure 7B). Since recall was a bigger limiting factor than precision in the *F*_1_ score, this resulted in prediction sets consisting of the union of predictions made by a combination of callers generally outperforming a single caller in terms of *F*_1_ as well as recall (Figure 4B). However, TEforest performed better (*F*_1_ score=0.852, recall=0.771, precision=0.954) than many more combinations of callers than other methods. While most combinations of callers that outperformed TEforest included TEforest, including the best combination of TEforest, TEMP, and PopoolationTE (*F*_1_ score=0.870, recall=0.813, precision=0.937), there were several combinations that did not, such as the third-best combination of TEMP, RetroSeq, and PopoolationTE (*F*_1_ score=0.867, recall=0.802, precision=0.942). While these results are informative as to how the union of different TE call sets may be used to improve performance, in light of the variation in accuracy that some methods show across read lengths and coverages, we caution readers that the relative performance of these different combinations may change across datasets.

### TEforest uses many features to make its predictions

We analyzed feature importances in TEforest’s non-reference model to identify the key features driving accurate TE detection and their relative contributions (Table 4), as measured by both permutation importance and impurity-based performance. Permutation importance measures the decrease in model performance when a feature’s values are randomly shuffled, reflecting the feature’s predictive power (Breiman 2001). Impurity-based importance, on the other hand, quantifies the average reduction in Gini impurity achieved by a feature during decision tree splits, highlighting its role in decision-making within the Random Forest model (Breiman 2002). The top ten features, ranked by permutation importance, were primarily related to discordantly mapping reads, split reads, and proper pairing of reads. In many cases the standard deviation or interquartile range (IQR) of a feature was more important for the model than central tendency metrics such as the mean or median. In general, features with high permutation importance had high impurity-importance, though the relative ordering of certain features differed for each method.

**Table 4:**
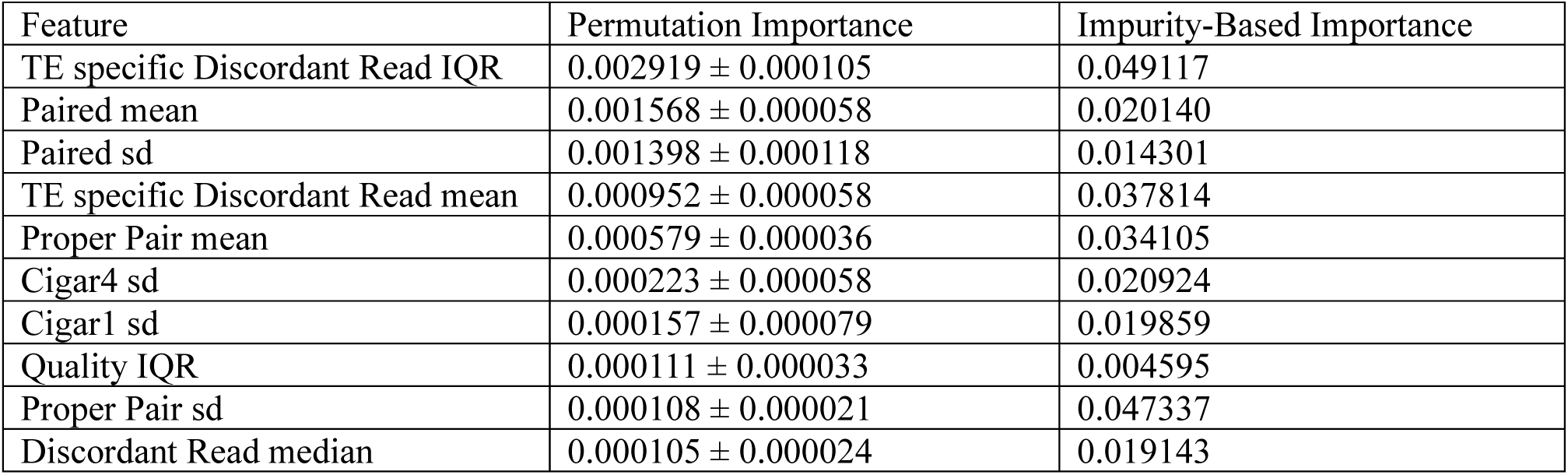
Feature importance metrics for the top non-reference model features. The table lists the top ten features ranked by permutation importance, which measures the decrease in model performance when feature values are randomly shuffled. Values are reported as the mean importance ± standard deviation across permutations. Impurity-based importance reflects the average reduction in Gini impurity achieved by the feature during decision tree splits in the Random Forest model.

One particularly informative feature was the TE-specific Discordant Read IQR, visualized in Figure 5. This feature exhibited clear stratification across model predictions: homozygotes had the highest values, heterozygotes showed intermediate values, and absences had the lowest values. However, overlap near the edges of these distributions highlighted the model’s reliance on additional features to maintain its high predictive performance. False positives had higher values for this feature compared to correctly predicted negatives, but they remained at the lower end of the range seen in true heterozygotes or homozygotes. Conversely, false negatives exhibited lower values than true positives but still exceeded those of true negatives. These patterns suggest that while TEforest excels at leveraging key features for prediction, the inclusion of complementary features is crucial for resolving edge cases and maintaining accuracy across complex scenarios.

**Figure 5:**
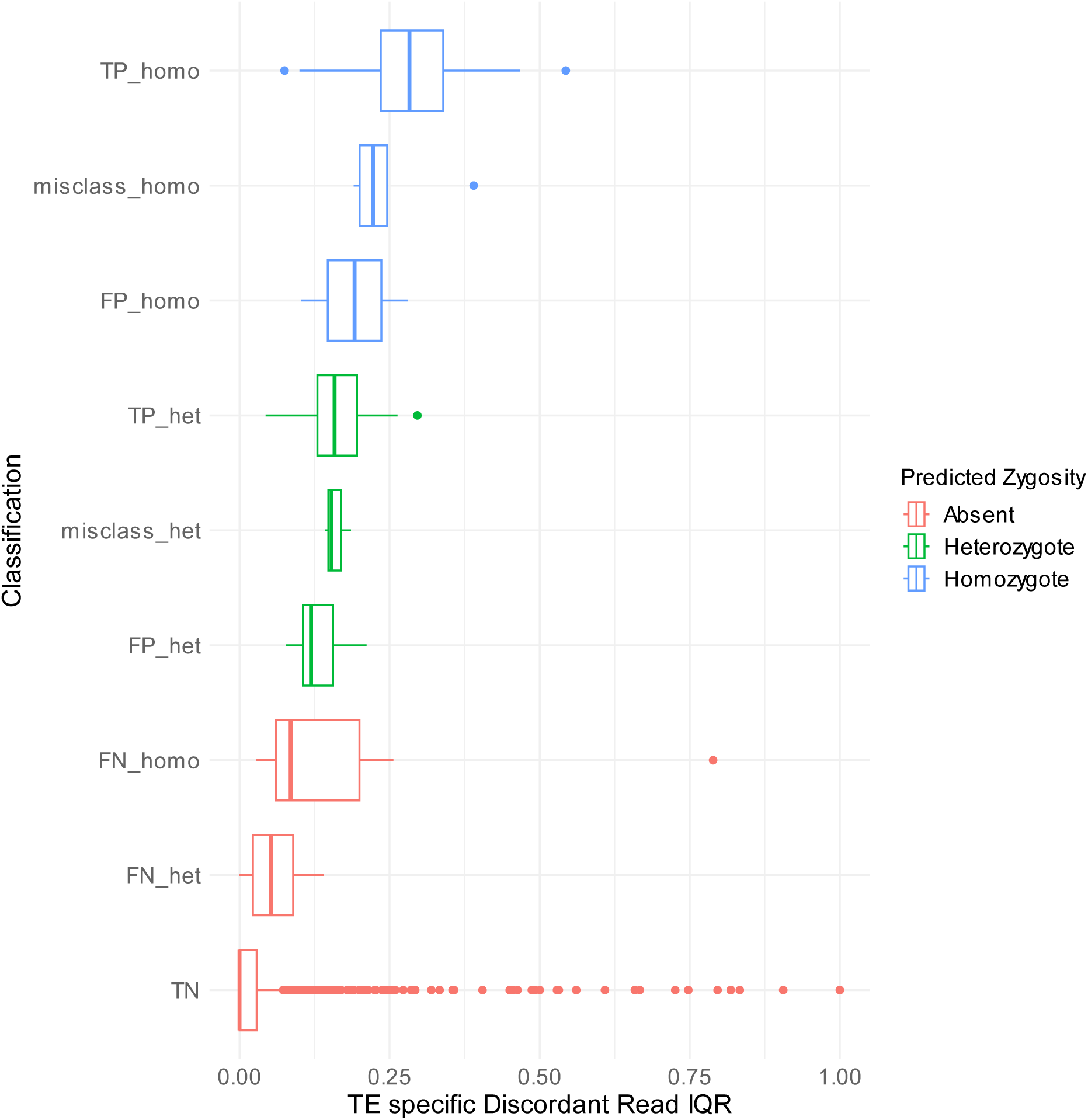
The distribution of the TE-specific discordant read interquartile range (IQR), the most important feature in the non-reference TE model, across various prediction categories, representing the relationship between true and predicted zygosity classifications. Categories include true positives (TP_homo, TP_het); true negatives (TN); candidate regions without true positives (FP_homo, FP_het); false negatives (FN_homo, FN_het); and misclassifications (misclass_homo, misclass_het), which involve errors such as misclassifying homozygotes or heterozygotes. Boxplots display the distribution of the values reported for each candidate region, with outliers shown as points. 11 outlier points >1 in the TN category were cropped out of this plot to improve the visualization.

### TEforest is robust to read lengths not in its training set, while TE callers show unexpected performance gains with shorter reads

To evaluate whether TEforest’s performance would decline when tested on read lengths not included in its training set, we conducted an analysis using trimmed reads from our synthetic genomes. Reads originally 151 and 125 bp in length were trimmed at the 3’ end down to 100 bp using *fastp* (Chen 2023), and the trimmed reads were downsampled to 30X genome coverage to enable a direct comparison with the previously tested 30X model.

For the reads that were trimmed from 151 bp down to 100 bp, TEforest encouragingly maintained an *F*_1_ score of 0.84 despite a 50% reduction in read length (SFigure 8). The recall for TEforest improved slightly from 0.75 to 0.76, while precision decreased to from 0.95 to 0.94.

These results suggest that TEforest’s feature vectors are resilient to variations in read length, preserving its ability to accurately identify TE insertions. Surprisingly, the performance of all other TE callers improved with these trimmed reads, with *F*_1_ scores increasing across the board. PopoolationTE2 exhibited the largest improvement, with its *F*_1_ score increasing from 0.32 to 0.46, while TEFLoN had the smallest improvement, from 0.55 to 0.57. Despite these gains, none of the callers outperformed TEforest. The improvements in other callers’ performance were primarily driven by higher recall, though this came at the cost of slightly reduced precision in most cases, with the exception of PopoolationTE2.

For the 125 bp reads, TEforest exhibited a slight decline in *F*_1_ score after trimming, dropping from 0.80 to 0.77 (SFigure 9). Recall decreased from 0.67 to 0.65, while precision declined from 0.97 to 0.96. In contrast, the performance of other TE callers improved even more dramatically with trimming than the 151 bp reads. RetroSeq showed the largest increase, with its *F*_1_ score rising from 0.33 to 0.68. Notably, TEMP2 surpassed TEforest’s performance, achieving an *F*_1_ score of 0.80, up from 0.76.

These unexpected increases in the performance of other TE callers may be attributed to their reliance on discordant and split reads that can be mapped to TE sequences. While split reads are a valuable resource for precisely detecting the breakpoints of TE insertions, it is more challenging to correctly map each piece of the split read compared to mapping a discordant read. When reads spanning a TE insertion are trimmed, the length of the effectively unsequenced portion of the fragment is increased, making it more likely that the two reads will be on either side of the breakpoint. This results in an increased number of discordant reads mapping to TE sequences (that is, non-TE mapping split reads become TE mapping discordant reads), likely enhancing the confidence of these tools in identifying TE insertions. In contrast, TEforest incorporates split-read information directly into its feature vectors, regardless of whether the read mapper correctly aligns the split piece back to the TE consensus sequence. This capability of TEforest is particularly desirable, as it allows researchers to utilize the full range of information present in their dataset without needing to manipulate or preprocess the data extensively to achieve optimal performance.

### Allele frequency prediction: Can we use TEforest for population genetics?

To characterize the impact of any class of mutation, it is essential not only to identify these mutations, but also to genotype them accurately enough to measure their frequency in a population. For example, the distribution of allele frequencies in a population, also called the site frequency spectrum (SFS) is a useful tool for understanding the population genetic forces governing a class of mutations (Williamson et al. 2005; Eyre-Walker et al. 2006; Keightley and Eyre-Walker 2007; Emerson et al. 2008). To accurately measure allele frequencies in a population, a TE caller will need to successfully detect and genotype the TE in all individuals carrying the allele (unless there are multiple detection/genotyping errors that happen to offset one another). For example, false positive predictions could skew the SFS towards rare mutations, potentially inflating estimates of the strength of natural selection acting against new mutations (Emerson et al. 2008; Johri et al. 2022).

To assess whether the genotyping accuracy of TE detection methods examined here is sufficient to produce accurate estimates of the SFS of TE insertions, we ran all previously tested TE detection methods in 13 genomes, sequenced with 54 bp reads, from the *Drosophila* synthetic population resource included in our truth dataset (Chakraborty et al. 2019; Rech et al. 2022). We assessed the allele frequency prediction accuracy for each non-reference insertion in the population by subtracting the true TE frequency from the number of true positive predictions (Figure 6A). Due to the tendency towards false negatives instead of false positives, all TE callers underpredicted the true TE frequency on average. TEMP and TEforest predicted frequencies more accurately than other callers, with TEMP2 and RetroSeq also performing well at this task, reflecting the previous benchmarking on single genomes. TEFLoN, PopoolationTE, and PopoolationTE2 struggled at this task due to many false negative predictions. When viewing the entire SFS for Roo (Figure 6B) and Copia (Figure 6C) insertions, which were the two TEs with the highest number of non-reference copies in this dataset, we found most callers predict the proportion of alleles in each frequency category fairly accurately. Perhaps in part because this dataset is comprised of a global sample of individuals rather than a local population, the vast majority of TE alleles were rare in the sample, limiting our ability to assess allele frequency prediction for high frequency alleles. However, combined with our previous benchmarking results, we find that TEforest is uniquely capable of both recovering TE insertions and obtaining accurate allele frequency estimates for downstream population genetic inference.

**Figure 6:**
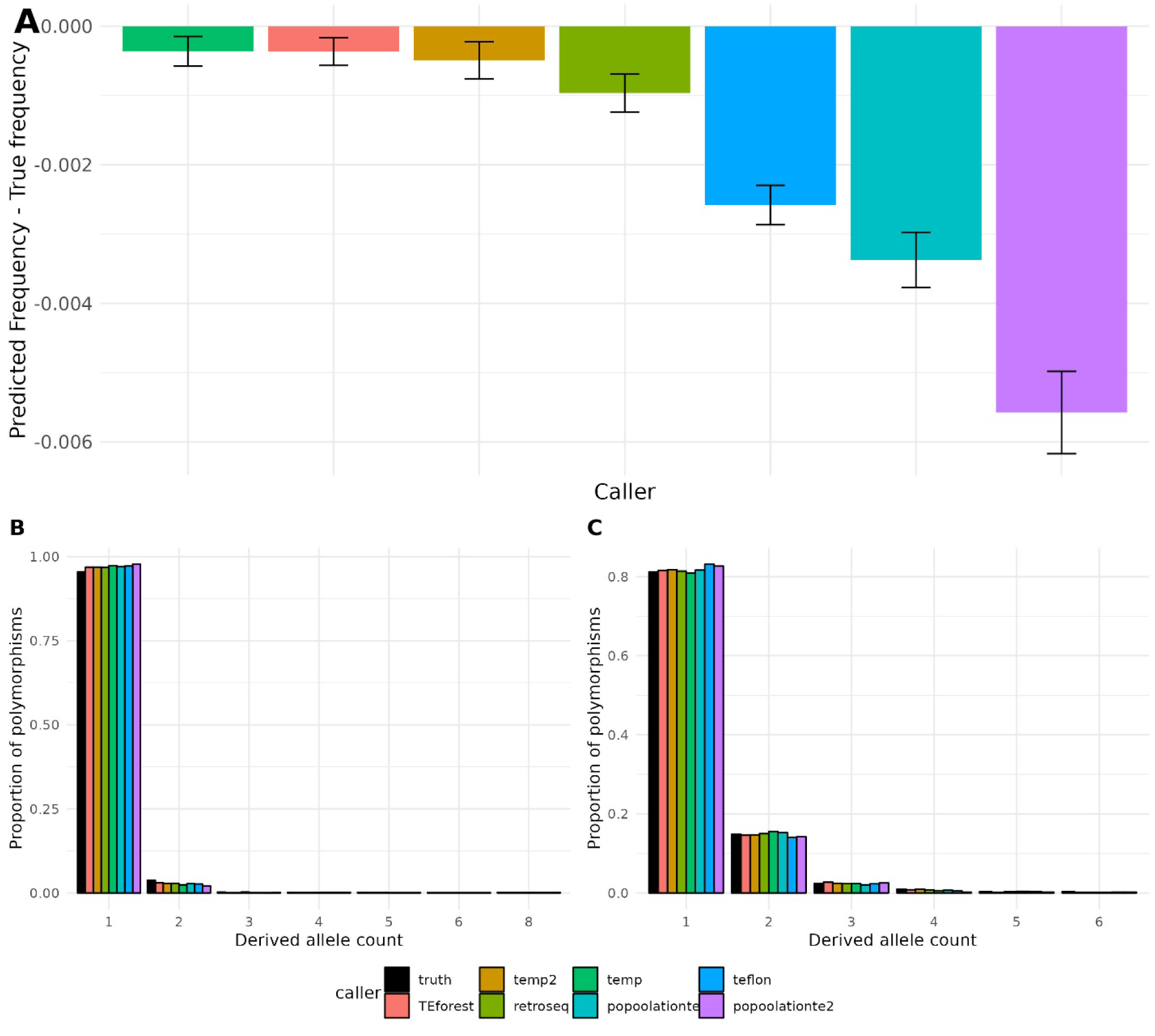
(**A**) Performance of TEforest compared to other TE callers at correctly predicting the allele frequencies of non-reference TE insertions in 13 fully homozygous individuals from the *Drosophila* synthetic population resource. The y axis represents the average difference in allele frequency predicted by each caller to the true allele frequency at that locus. Error bars represent the standard error for each caller. The site frequency spectrum of (**B**) roo and (**C**) jockey TE insertions, which had the highest number of TE insertions in these individuals.

### TEforest outperforms other methods in reference TE detection

We also trained a model to detect the presence or absence of TEs that are found in the reference genome, calculating the same feature vector as the non-reference model for each reference TE region. Since the locations of reference TEs are predefined by the user, the reference TE model does not require candidate region or breakpoint identification steps. Similar to the non-reference model, TEforest demonstrated superior accuracy in detecting the presence or absence of reference TEs compared to competing methods across the three tested read lengths and varying coverages (SFigure 10). Interestingly, this improvement in performance was due to a better balance between precision and recall compared to other models, rather than the increased recall observed in the non-reference model. With more informative reads available to detect reference TEs (there are often unique regions within each reference TE sequence where coverage changes depending on its presence or absence), TEforest, along with TEMP and TEMP2 (both of which use the same algorithm), maintained a high *F*_1_ score across all tested coverages. In contrast, TEFLoN and PopoolationTE2 had lower recall at 20X coverage and below.

## DISCUSSION

Long-read sequencing technologies are becoming increasingly affordable and useful for analyzing TEs (Chu et al. 2021; Han et al. 2022; Rech et al. 2022; Mohamed et al. 2023), but short-read sequencing remains the more accessible option for many researchers, especially for large-scale population studies. Moreover, the vast majority of population genomic datasets that are currently available to date consist of short-read sequences. To integrate recent advancements in long-read sequencing with existing population-scale short-read sequencing datasets, we developed TEforest—a machine learning tool for detecting and genotyping transposable element (TE) insertions using short-read data. By incorporating high-quality annotations from long-read *Drosophila melanogaster* genome assemblies in the training process, TEforest leverages the precision of long-read sequencing while taking advantage of the widespread availability of short-read data. Our comprehensive benchmarking, one of the most extensive real-data evaluations to date involving six different TE detection tools, demonstrates that TEforest not only outperforms existing methods but also maintains more stable performance across various read lengths and coverages. While other tools exhibit fluctuating performance depending on sequencing parameters, such as a surprising increase in performance with shorter read lengths, TEforest consistently delivers accurate detection and genotyping of TE insertions.

Our benchmarking assessments highlight the advantages of using real data over simulated data. While simulated datasets allow controlled testing of algorithms, currently do not capture the full complexity of real genomic data, including sequencing errors, structural variations near TE insertion breakpoints or within TE sequences, and the complexity of nested TE insertions.

Early testing on real data also shaped decisions made during the development of our algorithm, such as accounting for the fact that many TE insertion breakpoints have small gaps between the two breakpoints rather than the classic TSD signature (see the example insertion in Figure 1C). Such details might have been overlooked if we developed the algorithm using a simulated dataset.

In our study, we observed that in some cases PoPoolationTE and PoPoolationTE2, did not perform as well as other methods both in simulated datasets (also reported in Chen *et al*. (2023) and in our real datasets (see Vendrell-Mir *et al*. (2019) for a counter-example).

Conversely, TEMP2 and RetroSeq, which have shown strong performance in other studies, also performed reasonably well in our benchmarks.

However, our results also suggest that the combination of TE callers that performs the best on a given dataset will vary depending on the insert size, read length, sequencing coverage, and other species-specific parameters, and thus may be difficult to choose in practice without prior benchmarking. The benchmarks of previous studies have shown that TE detection tools often perform variably across different species due to differences in genome structure and TE content. For instance, tools like MELT (Gardner et al. 2017) originally developed for use in the human genome performed worse when applied to *Drosophila* genomes due to the mislabeling of related TE families, while TEMP2 and RetroSeq performed well in both contexts (Yu et al. 2021). Similarly, TEPID performed well when detecting simulated *Arabidopsis* insertions, but declined in performance when applied to *Drosophila* (Verneret et al. 2024). A potential solution to the variable performance of different methods depending on context is to use a combination of methods to make more confident predictions. Similar to Xu *et al*. (Xu et al. 2023), we find that certain combinations of TE detection methods can increase performance by increasing recall without a severe decline in precision. We also note that when evaluating different combinations of tools, we simply took the union of all TE insertions detected by a given set of callers, and it is possible that more complex rules for combining TE call sets from different methods could be more effective (e.g. requiring that all calls produced by a method with lower precision be recovered by at least one additional caller). In addition, one could optimize a model or model-ensemble to maximize precision at the cost of recall or vice-versa depending on the use case.

The tools XTea (Chu et al. 2021) and DeepMEI (Xu et al. 2023) also use machine learning in the context of non-reference TE detection. XTea employs random forest with a less comprehensive feature vector for short-read genotyping, while DeepMEI utilizes a convolutional neural network (CNN) for TE detection and genotyping. However, these tools are currently only available for use in human genomes and thus cannot be benchmarked against our method on this dataset.

The success of short-read TE detection approaches like TEforest is fundamentally dependent on the availability of high-quality reference genomes from which comprehensive TE annotations can be produced and for which short-read sequencing data can be obtained.

Thankfully, these resources are readily available in our tested species *D. melanogaster* (Kaminker et al. 2002; Rech et al. 2022). The performance of our approach of training on TE annotations and short-read mapping patterns from such assemblies would degrade when using poorly assembled or annotated reference genomes or species with an incomplete library of TE consensus sequences. Bearing this in mind, along with the often-unpredictable performance of TE detection tools when tested in a new species for which high quality training/benchmarking data are not available, we recommend careful evaluation of TE detection tools when analyzing TEs in non-model species, perhaps by simulating TE insertions as realistically as possible for testing (Chen et al. 2023; Verneret et al. 2024). Despite these challenges, we expect the performance and utility of short-read TE detection tools to increase as genome assemblies become more complete (Li and Durbin 2024) and tools to automatically curate TE consensus sequences and annotate them in a reference genome become more advanced (Baril et al. 2024; Orozco-Arias et al. 2024).

Our study, along with others (see Xu *et al*. (2023)), found that shorter, fragmented TE insertions are more challenging to detect than full-length insertions. As shorter TE copies are often inactivated or made reliant upon the transposition machinery encoded by other TE copies, developing extensions of TEforest that can predict whether insertions are full-length or fragmented using reads mapping to TE sequences could provide useful information about TE dynamics. While this approach could detect deletions on the edge of a TE due to changes in read-mapping patterns, its utility would be limited by the length of short reads, as they may lack coverage of the internal regions of TE insertions needed to detect internal deletions. For example, the longest paired-end reads used in this study would reach ∼400 bp into each side of the TE, while some TE families can exceed 10 kb in length. Another potential extension of TEforest involves detecting low-prevalence or somatic TE insertions, which are typically more challenging to detect due to the low number of informative reads per insertion (Yu et al. 2021); training data for such a method could be obtained by combining reads from genomes known to have different genotypes for a TE insertion to produce a pooled sample where the TE insertion is present at the desired prevalence. A machine learning algorithm like TEforest could be useful in distinguishing true insertions from false positives, provided that appropriate training data is available.

TEforest represents a significant step forward in the detection and genotyping of transposable element (TE) insertions from short-read sequencing data. By combining machine learning techniques with comprehensive feature extraction from read alignments, it consistently outperforms existing methods. As genomic resources and annotations continue to improve, approaches like TEforest—which leverage high quality ground-truth TE insertion datasets to enhance the utility of less costly sequencing data—will become increasingly robust. In turn, this will enable broader and more in-depth studies on the evolutionary and functional impact of TEs.

## ACKNOWLEDGEMENTS

We thank Marta Coronado Zamora and Josefa González for their assistance with the truth dataset. AD received support from the National Institute of General Medical Sciences of the National Institutes of Health under award numbers R35GM154969 and T32GM067553, and DRS received support from the National Institute of General Medical Sciences of the National Institutes of Health under award number R35GM138286.

## AUTHOR CONTRIBUTIONS

Austin Daigle: Conceptualization, Formal analysis, Software, Visualization, Writing—original draft, Writing—review & editing. Logan S. Whitehouse: Conceptualization, Formal analysis, Software, Writing—review & editing. Roy Zaho: Conceptualization, Resources, Writing— review & editing. Roy Zaho: Conceptualization, Resources, Writing—review & editing. Daniel R. Schrider: Conceptualization, Formal analysis, Visualization, Writing—review & editing, Funding acquisition.

## DATA AVAILABILITY

The code used to train and use TEforest, along with the trained models and code used for benchmarking, are available at https://github.com/SchriderLab/TEforest.git (figshare DOI: 10.6084/m9.figshare.28278599). The MCTE library of consensus TE sequences and annotations are avaliable at the DIGITAL.CSIC repository and can be accessed at http://doi.org/10.20350/DIGITALCSIC/13765 and http://doi.org/10.20350/DIGITALCSIC/13894, respectively. The short-read sequences used for this study are avaliable in the NCBI database under the BioProject accessions PRJNA559813 and SRP011971.

## REFERENCES

Adrion JR, Song MJ, Schrider DR, Hahn MW, Schaack S. 2017. Genome-wide estimates of transposable element insertion and deletion rates in *Drosophila melanogaster*. Genome Biol Evol. 9(5):1329–1340. doi:10.1093/gbe/evx050.

Baril T, Galbraith J, Hayward A. 2024. Earl Grey: A fully automated user-friendly transposable element annotation and analysis pipeline. Mol Biol Evol. 41(4):msae068. doi:10.1093/molbev/msae068.

Bourque G, Burns KH, Gehring M, Gorbunova V, Seluanov A, Hammell M, Imbeault M, Izsvák Z, Levin HL, Macfarlan TS, et al. 2018. Ten things you should know about transposable elements. Genome Biol. 19(1):199. doi:10.1186/s13059-018-1577-z.

Breiman L. 2001. Random forests. Mach Learn. 45(1):5–32. doi:10.1023/A:1010933404324.

Breiman L. 2002. Manual on setting up, using, and understanding random forests V3.1.

Casacuberta E, González J. 2013. The impact of transposable elements in environmental adaptation. Molecular Ecology. 22(6):1503–1517. doi:10.1111/mec.12170.

Chakraborty M, Emerson JJ, Macdonald SJ, Long AD. 2019. Structural variants exhibit widespread allelic heterogeneity and shape variation in complex traits. Nat Commun. 10(1):4872. doi:10.1038/s41467-019-12884-1.

Charlesworth B, Langley CH. 1989. The population genetics of *Drosophila* transposable elements. Ann Rev Genet. 23(Volume 23,):251–287. doi:10.1146/annurev.ge.23.120189.001343.

Chen J, Basting PJ, Han S, Garfinkel DJ, Bergman CM. 2023. Reproducible evaluation of transposable element detectors with McClintock 2 guides accurate inference of *Ty* insertion patterns in yeast. Mob DNA. 14(1):8. doi:10.1186/s13100-023-00296-4.

Chen S. 2023. Ultrafast one-pass FASTQ data preprocessing, quality control, and deduplication using fastp. iMeta. 2(2):e107. doi:10.1002/imt2.107.

Chu C, Borges-Monroy R, Viswanadham VV, Lee S, Li H, Lee EA, Park PJ. 2021. Comprehensive identification of transposable element insertions using multiple sequencing technologies. Nat Commun. 12(1):3836. doi:10.1038/s41467-021-24041-8.

Craig NL. 2007. Mobile DNA: an Introduction. In: Mobile DNA II. John Wiley & Sons, Ltd. p. 1–11. https://onlinelibrary.wiley.com/doi/abs/10.1128/9781555817954.ch1.

Danecek P, Bonfield JK, Liddle J, Marshall J, Ohan V, Pollard MO, Whitwham A, Keane T, McCarthy SA, Davies RM, et al. 2021. Twelve years of SAMtools and BCFtools. GigaScience. 10(2):giab008. doi:10.1093/gigascience/giab008.

Drongitis D, Aniello F, Fucci L, Donizetti A. 2019. Roles of transposable elements in the different layers of gene expression regulation. Int J Mol Sci. 20(22):5755. doi:10.3390/ijms20225755.

Emerson JJ, Cardoso-Moreira M, Borevitz JO, Long M. 2008. Natural selection shapes genome-wide patterns of copy-number polymorphism in *Drosophila melanogaster*. Science. 320(5883):1629–1631. doi:10.1126/science.1158078.

Eyre-Walker A, Woolfit M, Phelps T. 2006. The distribution of fitness effects of new deleterious amino acid mutations in humans. Genetics. 173(2):891–900. doi:10.1534/genetics.106.057570.

Feschotte C. 2008. Transposable elements and the evolution of regulatory networks. Nat Rev Genet. 9(5):397–405. doi:10.1038/nrg2337.

Finnegan DJ. 1992. Transposable elements. Curr Opin Genet Dev. 2(6):861–867. doi:10.1016/S0959-437X(05)80108-X.

Gardner EJ, Lam VK, Harris DN, Chuang NT, Scott EC, Pittard WS, Mills RE, Consortium T 1000 GP, Devine SE. 2017. The Mobile Element Locator Tool (MELT): population-scale mobile element discovery and biology. Genome Res. 27(11):1916–1929. doi:10.1101/gr.218032.116.

Han S, Dias GB, Basting PJ, Viswanatha R, Perrimon N, Bergman CM. 2022. Local assembly of long reads enables phylogenomics of transposable elements in a polyploid cell line. Nucleic Acids Research. 50(21):e124. doi:10.1093/nar/gkac794.

Hill T, Unckless RL. 2019. A deep learning approach for detecting copy number variation in next-generation sequencing data. G3. 9(11):3575–3582. doi:10.1534/g3.119.400596.

Hoyt SJ, Storer JM, Hartley GA, Grady PGS, Gershman A, de Lima LG, Limouse C, Halabian R, Wojenski L, Rodriguez M, et al. 2022. From telomere to telomere: The transcriptional and epigenetic state of human repeat elements. Science. 376(6588):eabk3112. doi:10.1126/science.abk3112.

Johri P, Aquadro CF, Beaumont M, Charlesworth B, Excoffier L, Eyre-Walker A, Keightley PD, Lynch M, McVean G, Payseur BA, et al. 2022. Recommendations for improving statistical inference in population genomics. PLoS Biol. 20(5):e3001669. doi:10.1371/journal.pbio.3001669.

Kaminker JS, Bergman CM, Kronmiller B, Carlson J, Svirskas R, Patel S, Frise E, Wheeler DA, Lewis SE, Rubin GM, et al. 2002. The transposable elements of the *Drosophila melanogaster* euchromatin: a genomics perspective. Genome Biol. 3(12):research0084.1. doi:10.1186/gb-2002-3-12-research0084.

Kapitonov VV, Jurka J. 2003. Molecular paleontology of transposable elements in the *Drosophila melanogaster* genome. Proc Natl Acad Sci USA. 100(11):6569–6574. doi:10.1073/pnas.0732024100.

Keane TM, Wong K, Adams DJ. 2013. RetroSeq: transposable element discovery from next-generation sequencing data. Bioinformatics. 29(3):389–90. doi:10.1093/bioinformatics/bts697.

Keightley PD, Eyre-Walker A. 2007. Joint inference of the distribution of fitness effects of deleterious mutations and population demography based on nucleotide polymorphism frequencies. Genetics. 177(4):2251–2261. doi:10.1534/genetics.107.080663.

Kirov I, Merkulov P, Dudnikov M, Polkhovskaya E, Komakhin RA, Konstantinov Z, Gvaramiya S, Ermolaev A, Kudryavtseva N, Gilyok M, et al. 2021. Transposons hidden in *Arabidopsis thaliana* genome assembly gaps and mobilization of non-autonomous LTR retrotransposons unravelled by Nanotei pipeline. Plants. 10(12):2681. doi:10.3390/plants10122681.

Kofler R, Betancourt AJ, Schlötterer C. 2012. Sequencing of pooled DNA samples (Pool-Seq) uncovers complex dynamics of transposable element insertions in *Drosophila melanogaster*. PLoS Genet. 8(1):e1002487. doi:10.1371/journal.pgen.1002487.

Kofler R, Gómez-Sánchez D, Schlötterer C. 2016. PoPoolationTE2: Comparative population genomics of transposable elements using pool-seq. Mol Biol Evol. 33(10):2759–2764. doi:10.1093/molbev/msw137.

Lawrence M, Huber W, Pagès H, Aboyoun P, Carlson M, Gentleman R, Morgan MT, Carey VJ. 2013. Software for computing and annotating genomic ranges. PLoS Comput Biol. 9(8):e1003118. doi:10.1371/journal.pcbi.1003118.

Lee YCG. 2015. The role of piRNA-mediated epigenetic silencing in the population dynamics of transposable elements in *Drosophila melanogaster*. PLoS Genet. 11(6):e1005269. doi:10.1371/journal.pgen.1005269.

Li H, Durbin R. 2009. Fast and accurate short read alignment with Burrows–Wheeler transform. Bioinformatics. 25(14):1754–1760. doi:10.1093/bioinformatics/btp324.

Li H, Durbin R. 2024. Genome assembly in the telomere-to-telomere era. Nat Rev Genet. 25(9):658–670. doi:10.1038/s41576-024-00718-w.

Linheiro RS, Bergman CM. 2012. Whole genome resequencing reveals natural target site preferences of transposable elements in *Drosophila melanogaster*. PLoS One. 7(2):e30008. doi:10.1371/journal.pone.0030008.

Lynch M. 2007. The origins of genome architecture. Sunderland, Mass: Sinauer Associates.

Makałowski W, Gotea V, Pande A, Makałowska I. 2019. Transposable elements: classification, identification, and their use as a tool for comparative genomics. In: Anisimova M, editor. Evolutionary Genomics: Statistical and Computational Methods. New York, NY: Springer. p. 177–207. 10.1007/978-1-4939-9074-0_6.

Mohamed M, Sabot F, Varoqui M, Mugat B, Audouin K, Pélisson A, Fiston-Lavier A-S, Chambeyron S. 2023. TrEMOLO: accurate transposable element allele frequency estimation using long-read sequencing data combining assembly and mapping-based approaches. Genome Biol. 24(1):63. doi:10.1186/s13059-023-02911-2.

Montgomery E, Charlesworth B, Langley CH. 1987. A test for the role of natural selection in the stabilization of transposable element copy number in a population of *Drosophila melanogaster*. Genet Res. 49(1):31–41. doi:10.1017/S0016672300026707.

Montgomery EA, Huang SM, Langley CH, Judd BH. 1991. Chromosome rearrangement by ectopic recombination in *Drosophila melanogaster*: genome structure and evolution. Genetics. 129(4):1085–1098. doi:10.1093/genetics/129.4.1085.

Nelson MG, Linheiro RS, Bergman CM. 2017. McClintock: an integrated pipeline for detecting transposable element insertions in whole-genome shotgun sequencing data. G3. 7(8):2763. doi:10.1534/g3.117.043893.

Orozco-Arias S, Sierra P, Durbin R, González J. 2024 Dec 9. MCHelper automatically curates transposable element libraries across eukaryotic species. Genome Res. doi:10.1101/gr.278821.123. https://genome.cshlp.org/content/early/2024/12/09/gr.278821.123.

Pedregosa F, Varoquaux G, Gramfort A, Michel V, Thirion B, Grisel O, Blondel M, Prettenhofer P, Weiss R, Dubourg V, et al. 2011. Scikit-learn: Machine learning in python. JMLR. 12(85):2825–2830.

Rech GE, Radío S, Guirao-Rico S, Aguilera L, Horvath V, Green L, Lindstadt H, Jamilloux V, Quesneville H, González J. 2022. Population-scale long-read sequencing uncovers transposable elements associated with gene expression variation and adaptive signatures in Drosophila. Nat Commun. 13(1):1948. doi:10.1038/s41467-022-29518-8.

Rishishwar L, Mariño-Ramírez L, Jordan IK. 2017. Benchmarking computational tools for polymorphic transposable element detection. Briefings in Bioinformatics. 18(6):908–918. doi:10.1093/bib/bbw072.

Schrader L, Schmitz J. 2019. The impact of transposable elements in adaptive evolution. Mol Ecol. 28(6):1537–1549. doi:10.1111/mec.14794.

Shen W, Sipos B, Zhao L. 2024. SeqKit2: A Swiss army knife for sequence and alignment processing. iMeta. 3(3):e191. doi:10.1002/imt2.191.

Singh ND, Petrov DA. 2004. Rapid sequence turnover at an intergenic locus in *Drosophila*. Mol Biol Evol. 21(4):670–680. doi:10.1093/molbev/msh060.

Vasimuddin Md, Misra S, Li H, Aluru S. 2019. Efficient architecture-aware acceleration of BWA-MEM for multicore systems. In: 2019 IEEE International Parallel and Distributed Processing Symposium (IPDPS). p. 314–324. https://ieeexplore.ieee.org/document/8820962.

Vendrell-Mir P, Barteri F, Merenciano M, González J, Casacuberta JM, Castanera R. 2019. A benchmark of transposon insertion detection tools using real data. Mobile DNA. 10(1):53. doi:10.1186/s13100-019-0197-9.

Verneret M, Le VA, Faraut T, Turpin J, Lerat E. 2024. Particular sequence characteristics induce bias in the detection of polymorphic transposable element insertions.:2024.09.25.614865. doi:10.1101/2024.09.25.614865. https://www.biorxiv.org/content/10.1101/2024.09.25.614865v2.

Wells JN, Feschotte C. 2020. A field guide to eukaryotic transposable elements. Annu Rev Genet. 54(1):539–561. doi:10.1146/annurev-genet-040620-022145.

Williamson SH, Hernandez R, Fledel-Alon A, Zhu L, Nielsen R, Bustamante CD. 2005. Simultaneous inference of selection and population growth from patterns of variation in the human genome. Proc Natl Acad Sci USA. 102(22):7882–7887. doi:10.1073/pnas.0502300102.

Xu X, Huang Y, Wang X, Cheng J, Yuan H, Bu F. 2023. Identification of mobile element insertion from whole genome sequencing data using deep neural network model.:2023.03.07.531451. doi:10.1101/2023.03.07.531451. https://www.biorxiv.org/content/10.1101/2023.03.07.531451v1.

Yu T, Huang X, Dou S, Tang X, Luo S, Theurkauf WE, Lu J, Weng Z. 2021. A benchmark and an algorithm for detecting germline transposon insertions and measuring de novo transposon insertion frequencies. Nucleic Acids Res. 49(8):e44. doi:10.1093/nar/gkab010.

Zhuang J, Wang J, Theurkauf W, Weng Z. 2014. TEMP: a computational method for analyzing transposable element polymorphism in populations. Nucleic Acids Res. 42(11):6826–6838. doi:10.1093/nar/gku323.

