## Supplemental File for "Leveraging long-read assemblies and machine learning to enhance short-read transposable element detection and genotyping"

### SUPPLEMENTARY FIGURES AND TABLES

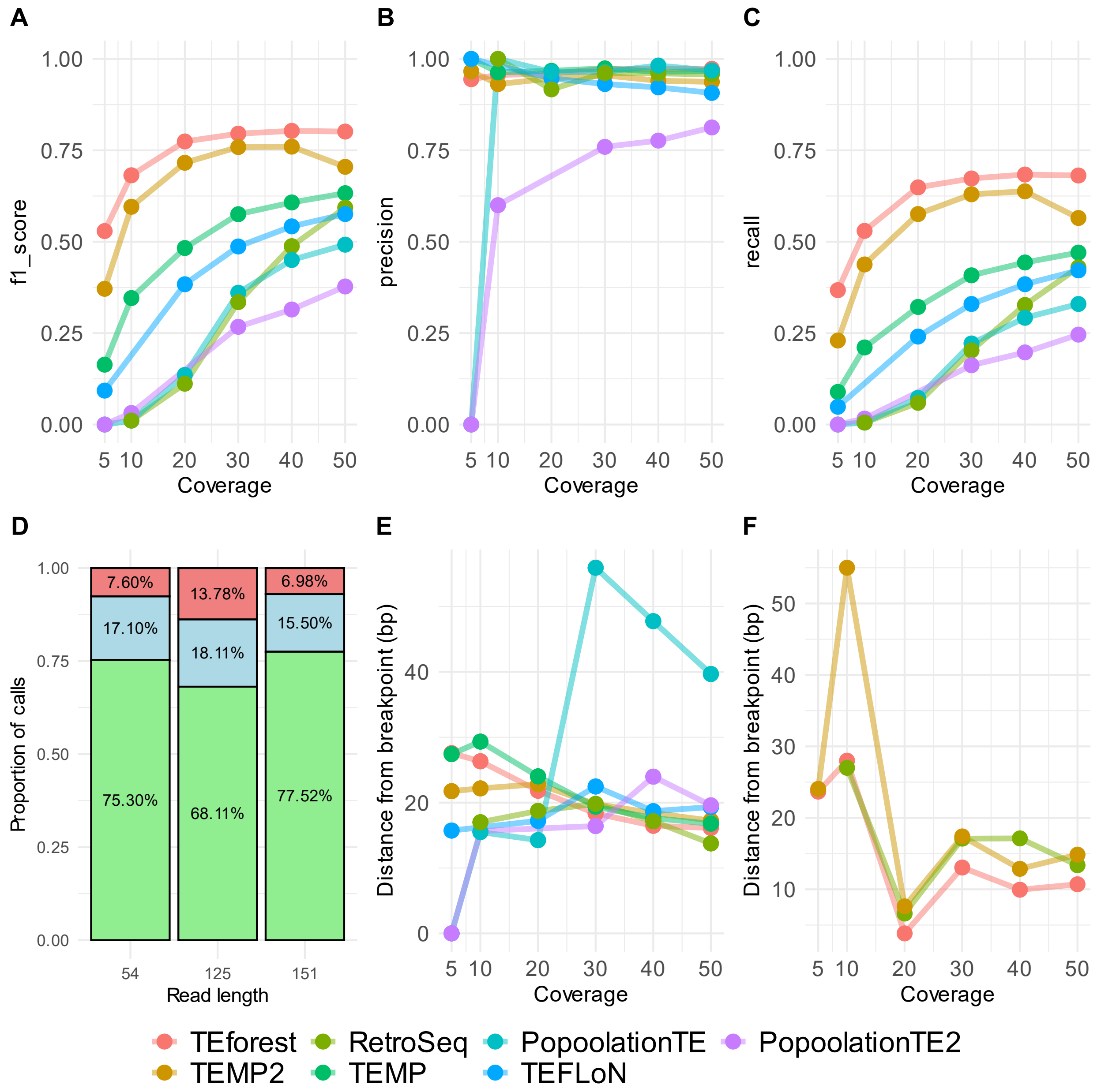


**Supplementary Figure 1:** The performance of TEforest compared to other short-read TE callers at detecting nonreference TE insertions annotated by long read assemblies of *D. melanogaster* strains. Short reads used for detection of TEs were 125 bp with ~208 bp insert sizes. True positives were defined as predicting a TE insertion of the correct TE family with any prevalence within 500 bp of the true insertion site. Performance was quantified with (**A**) F1 scores, (**B**) precision, and (**C**) recall. The proportions of TEs that were successfully detected by TEforest, lost in candidate region detection stage, or misclassified by the random forest model for each read length are shown in Panel **D**. The mean breakpoint accuracy of TE callers for (**E**) all true positive calls or (**F**) true positive calls shared by TEforest, RetroSeq and TEMP2 was quantified by finding the distance between the center of true positive breakpoint ranges predicted by the TE callers and the breakpoint in the truth dataset.

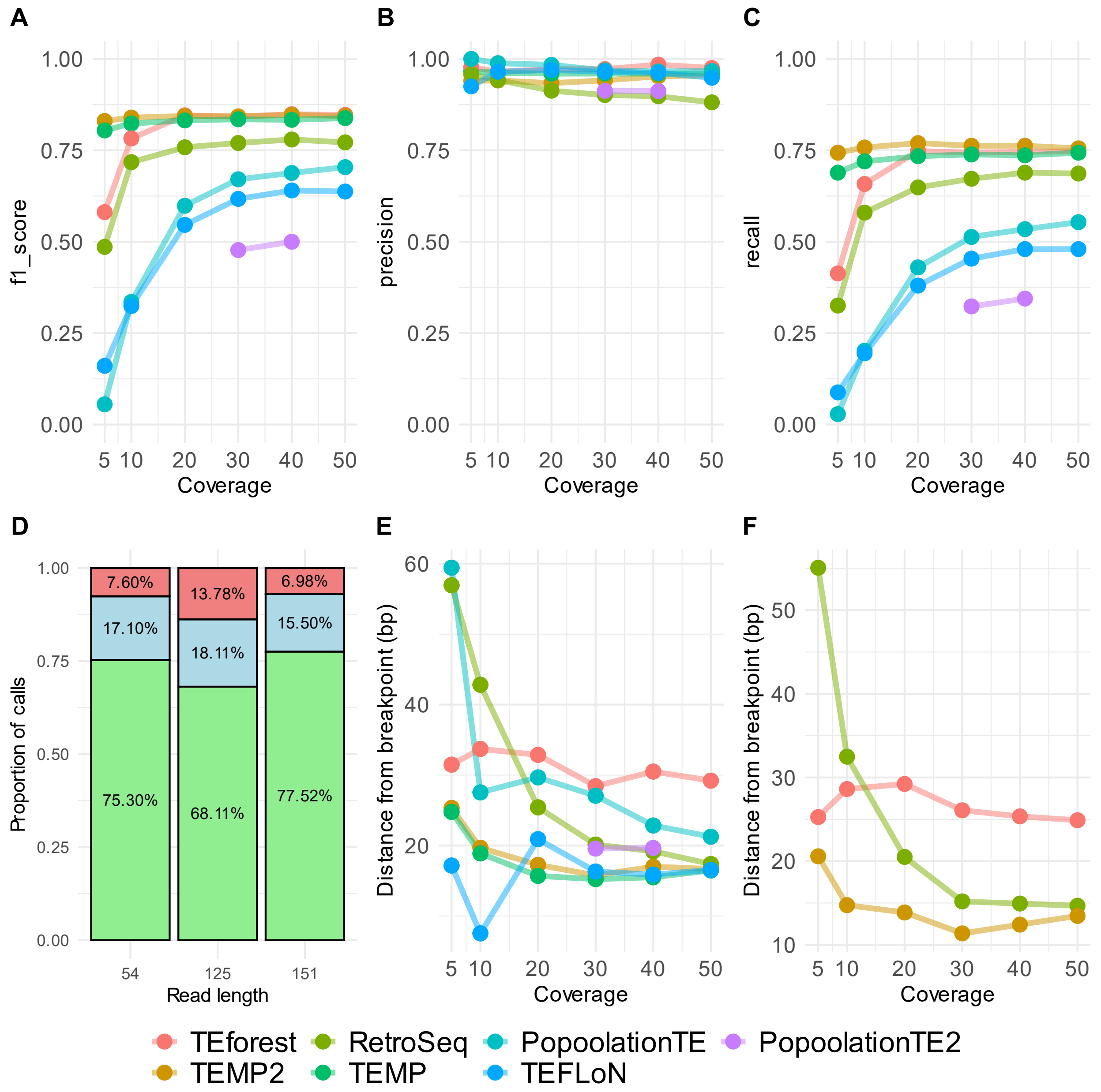


**Supplementary Figure 2:** The performance of TEforest compared to other short-read TE callers at detecting nonreference TE insertions annotated by long read assemblies of *D. melanogaster* strains. Short reads used for detection of TEs were 54 bp with ~287 bp insert sizes. True positives were defined as predicting a TE insertion of the correct TE family with any prevalence within 500 bp of the true insertion site. Performance was quantified with (**A**) F1 scores, (**B**) precision, and (**C**) recall. The proportions of TEs that were successfully detected by TEforest, lost in candidate region detection stage, or misclassified by the random forest model for each read length are shown in Panel **D**. The mean breakpoint accuracy of TE callers for (**E**) all true positive calls or (**F**) true positive calls shared by TEforest, RetroSeq and TEMP2 was quantified by finding the distance between the center of true positive breakpoint ranges predicted by the TE callers and the breakpoint in the truth dataset.

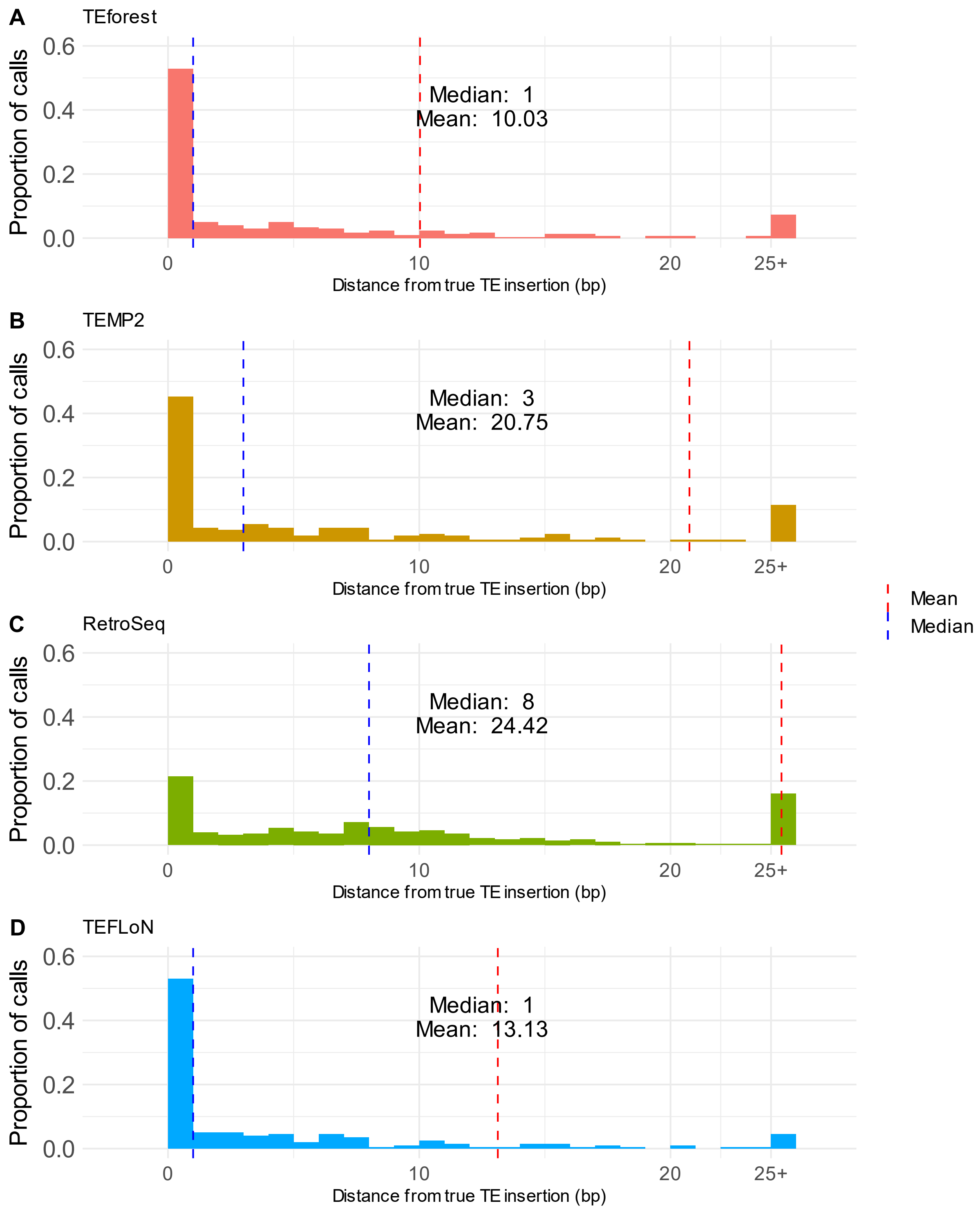


**Supplementary Figure 3:** The normalized distribution of breakpoint accuracy for all true positive calls for the 154 bp dataset from **(A)** TEforest, **(B)** TEMP2, **(C)** RetroSeq, and **(D)** TEFLoN was quantified by finding the distance between the center of true positive breakpoint ranges predicted by the TE callers and the breakpoint in the truth dataset. The last bin represents the proportion of calls >=25 bp away from the annotated breakpoint.

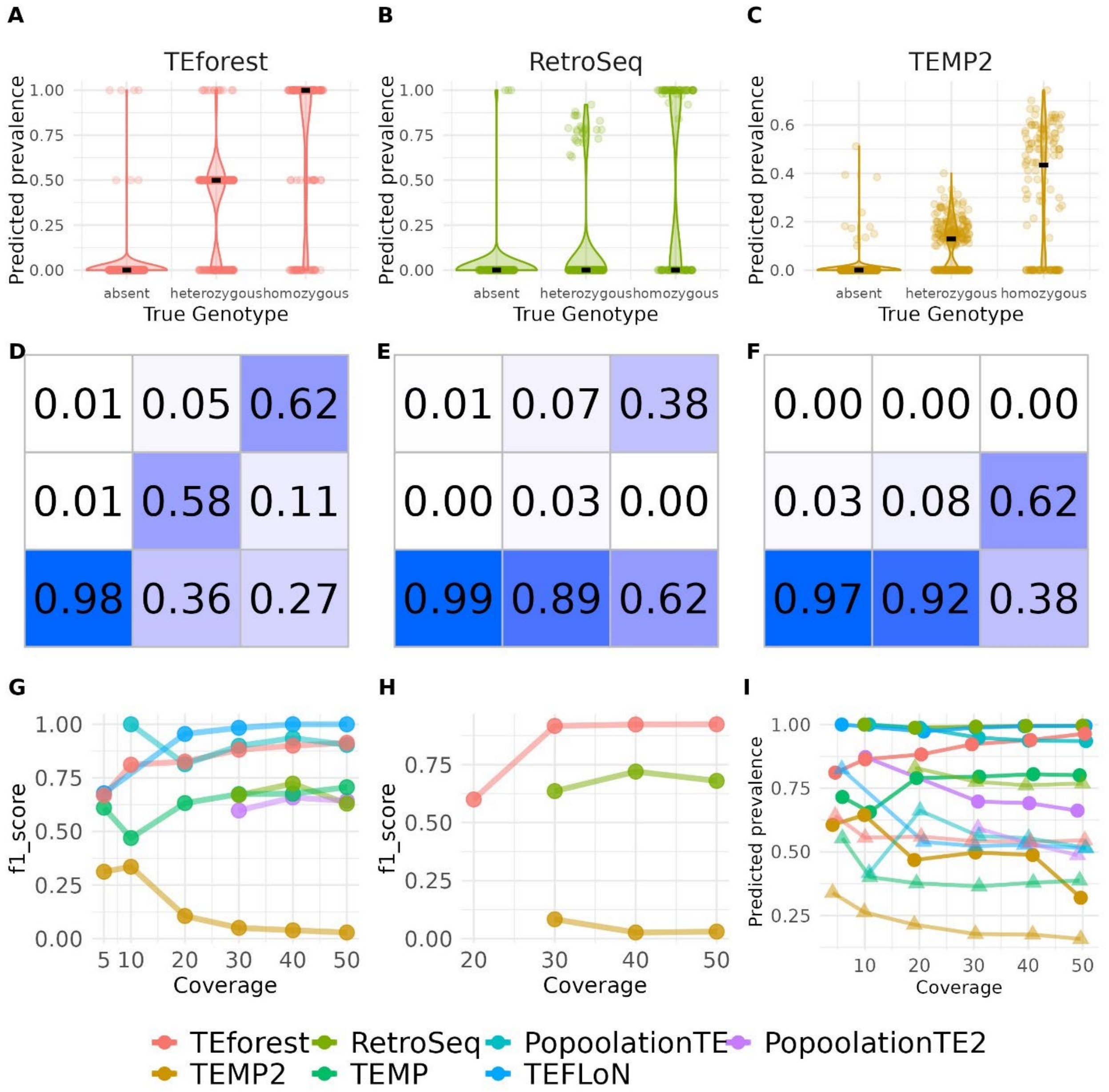


**Supplementary Figure 4:** The performance of TEforest compared to other short-read TE callers at genotyping nonreference TE insertions. Short reads used for detection of TEs were 125 bp with ~208 bp insert sizes. Frequency predictions for insertions that were homozygous, heterozygous, or absent were calculated for (**A**) TEforest, (**B**) RetroSeq, and (**C**) TEMP2, where true positive absences refer to candidate regions identified by TEforest with no TE insertion. The median prevalence is shown as a black line. (**D**-**F**) The accuracy of these frequency predictions is quantified in confusion matrices, where heterozygous predictions are defined as frequency predictions between 0.25 and 0.75 and homozygous predictions are above 0.75. (**G**) The genotyping F1 score for all true positive calls or (**H**) true positive calls shared by TEforest, RetroSeq and TEMP2 was quantified using true positive predictions of each caller. The mean for the predicted prevalence of true positive homozygous (circles) and heterozygous (triangles) insertions are shown in panel **I**, calculated using only the prevalences of true positive predictions of each caller (false positives are excluded).

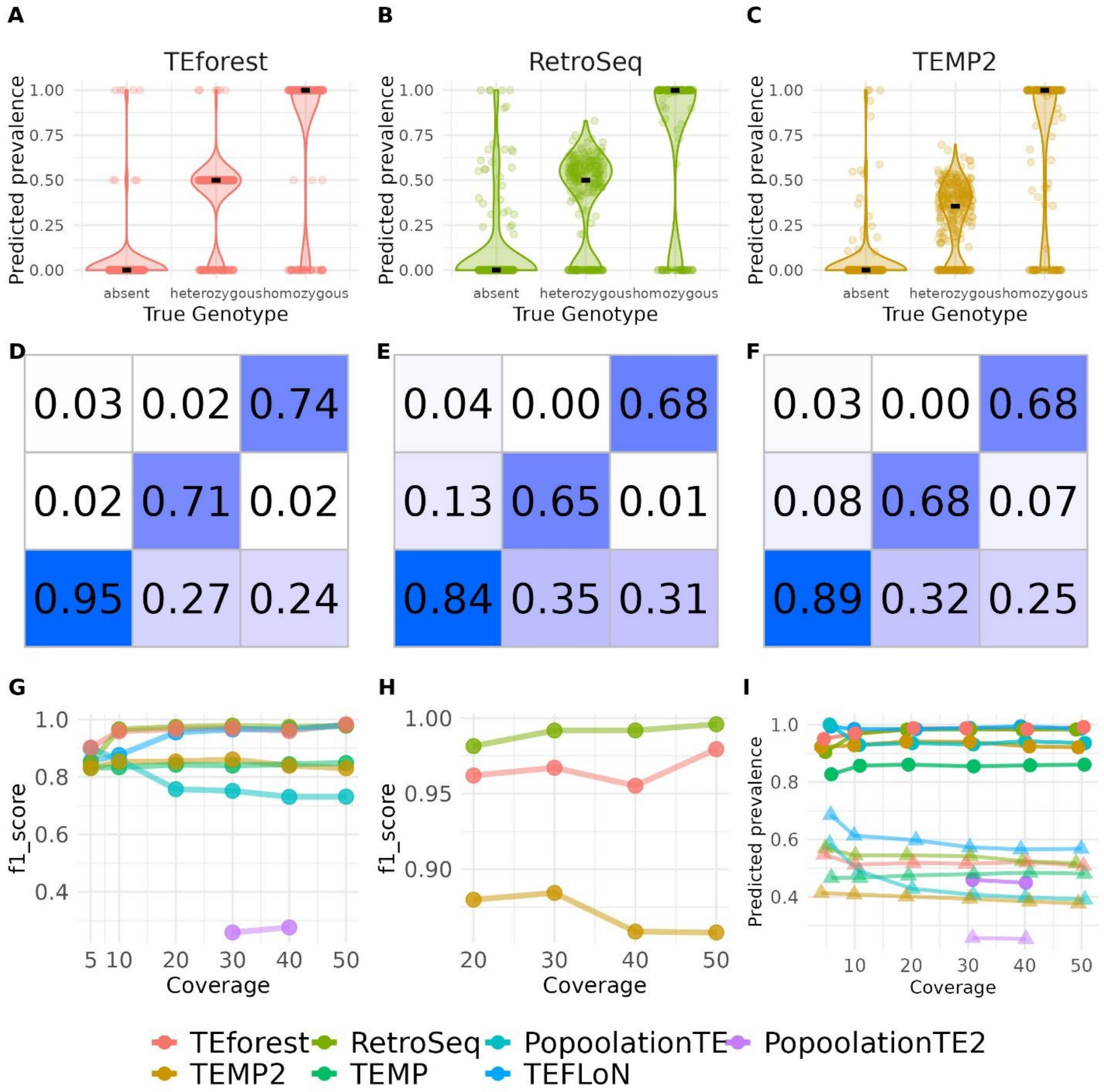


**Supplementary Figure 5:** The performance of TEforest compared to other short-read TE callers at genotyping nonreference TE insertions. Short reads used for detection of TEs were 54 bp with ~287 bp insert sizes. Frequency predictions for insertions that were homozygous, heterozygous, or absent were calculated for (**A**) TEforest, (**B**) RetroSeq, and (**C**) TEMP2, where true positive absences refer to candidate regions identified by TEforest with no TE insertion. The median prevalence is shown as a black line. (**D**-**F**) The accuracy of these frequency predictions is quantified in confusion matrices, where heterozygous predictions are defined as frequency predictions between 0.25 and 0.75 and homozygous predictions are above 0.75. (**G**) The genotyping F1 score for all true positive calls or (**H**) true positive calls shared by TEforest, RetroSeq and TEMP2 was quantified using true positive predictions of each caller. The mean for the predicted prevalence of true positive homozygous (circles) and heterozygous (triangles) insertions are shown in panel **I**, calculated using only the prevalences of true positive predictions of each caller (false positives are excluded).

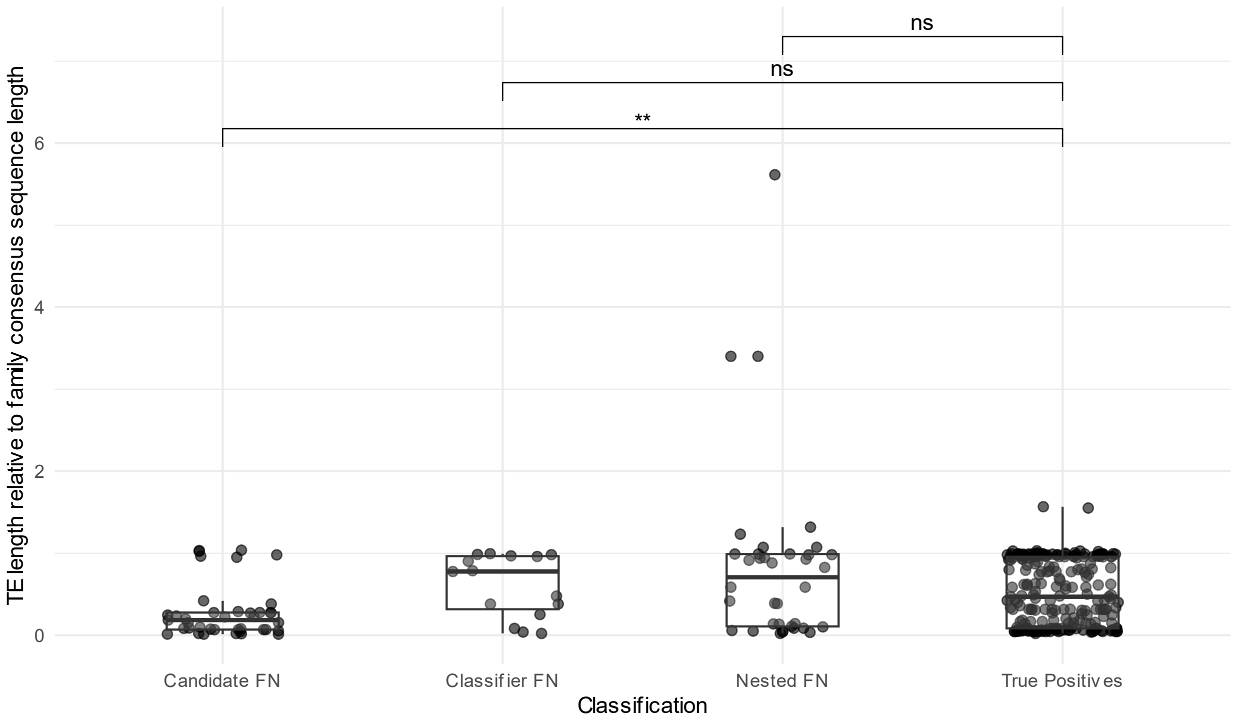


**Supplementary Figure 6:** Ratio of TE sequence length to the consensus length of that TE family of non-reference TEs insertions that were in the genome with 151 bp with ~436 bp insert sizes, divided into classes depending on whether they were unnested and not detected in the candidate region identification stage (Candidate FN), unnested and mislabeled as absences by the random forest classifier (Classifier FN), nested and lost in either the candidate regions or classifier step (Nested FN), or successfully detected by TEforest (True Positives). Overhead bars represent the results of pairwise Wilcox rank-sum tests.

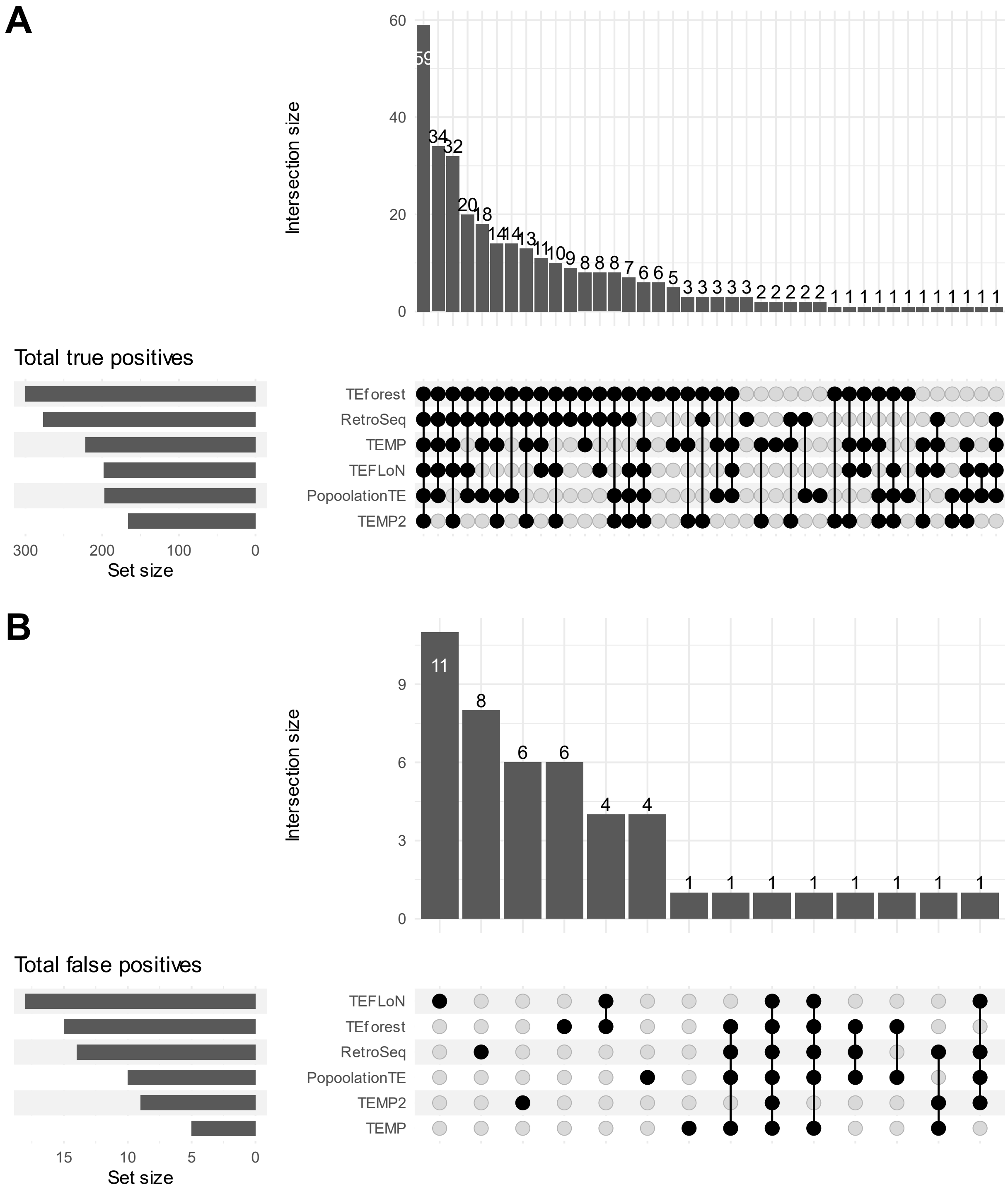


**Supplementary Figure 7: (A)** Upset plot representing the number of unique true positives for different combinations of TE callers. True positives were defined as a prediction made within 500 bp of an annotation within the truth dataset. Nested TEs from the same family were only counted once. **(B)** Upset plot representing the number of unique false positives for different combinations of TE callers.

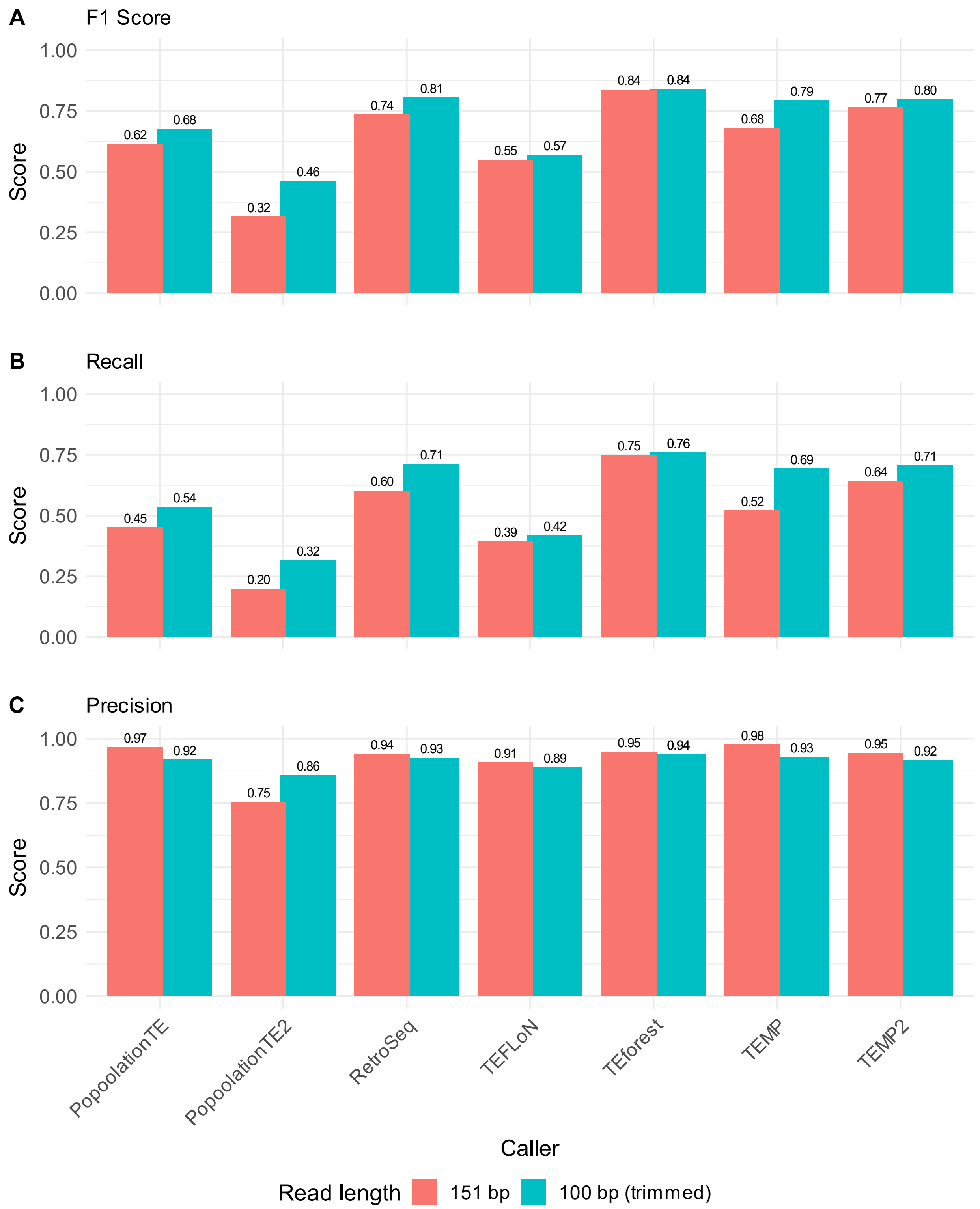


**Supplementary Figure 8:** Effects of trimming reads on the (**A**) F1 score, (**B**) recall, and (**C**) precision of TEforest and other short-read TE callers at detecting non-reference TE insertions. Reads were originally 151 bp with ~436 bp insert sizes, then shortened to 100 bp with the same insert size by trimming at the 3' end of the reads. Both datasets were downsampled to 30X coverage.

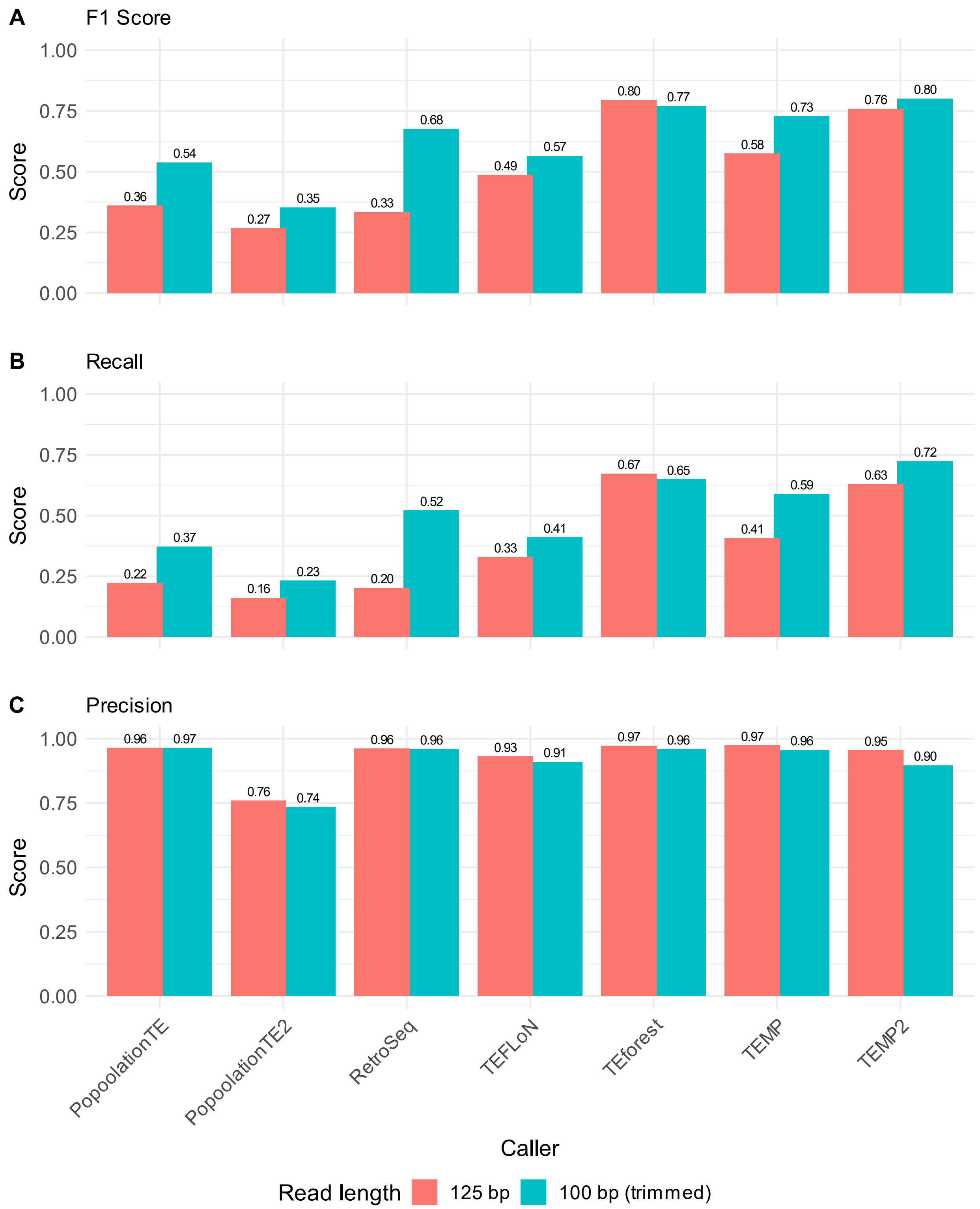


**Supplementary Figure 9:** Effects of trimming reads on the (**A**) F1 score, (**B**) recall, and (**C**) precision of TEforest and other short-read TE callers at detecting non-reference TE insertions. Reads were originally 125 bp with ~208 bp insert sizes, then shortened to 100 bp with the same insert size by trimming at the 3' end of the reads. Both datasets were downsampled to 30X coverage.

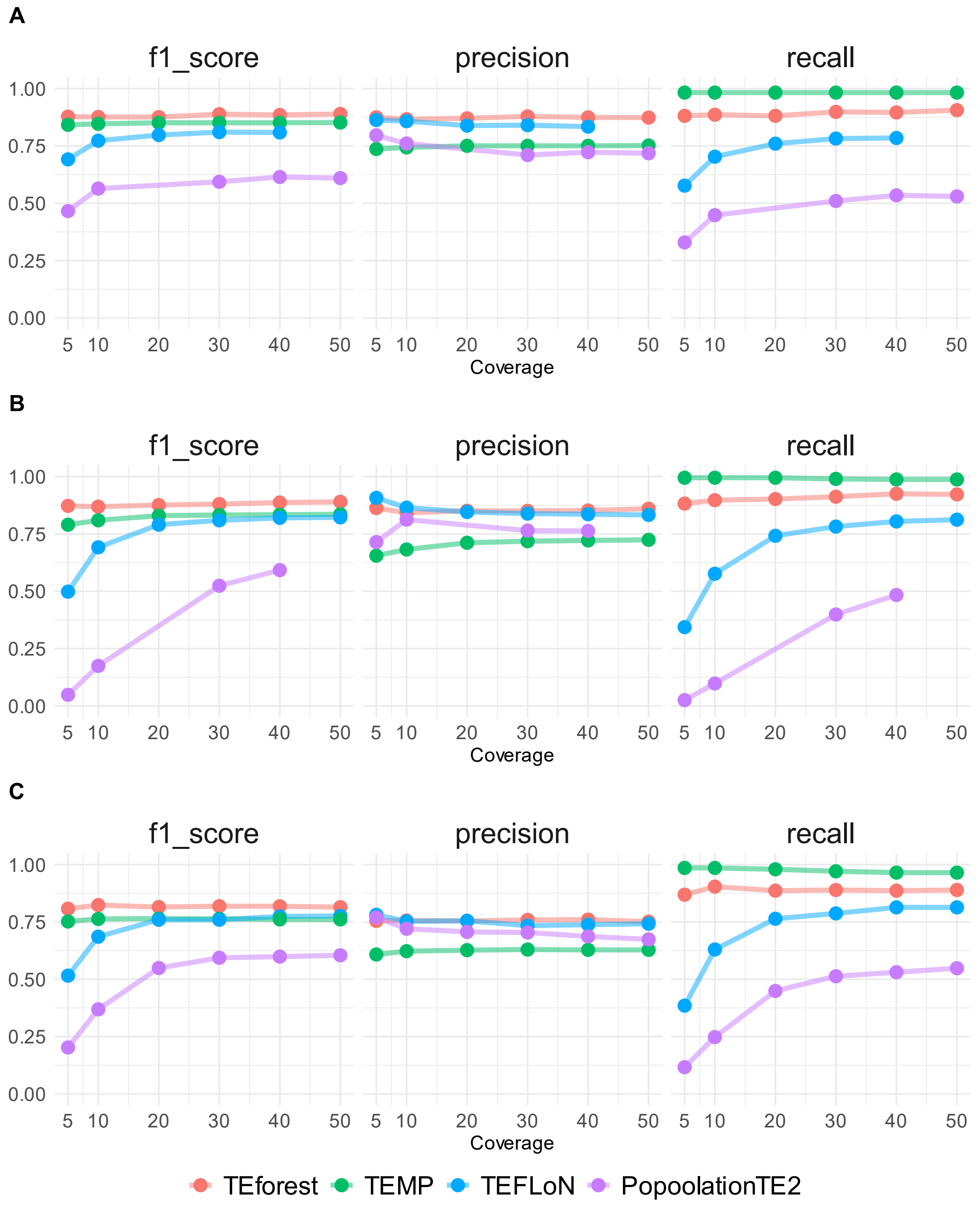


**Supplementary Figure 10:** The performance of TEforest compared to other short-read TE callers at detecting reference TE insertions annotated by long read assemblies of *D. melanogaster* strains. Short reads used for detection of TEs were (**A**) 54 bp with ~436 bp insert sizes, (**B**) 125 bp with ~208 bp insert sizes, and (**C**) 151 bp with ~208 bp insert sizes. Performance was quantified with F1 scores, precision, and recall.

**Supplementary Table 1:** TE family specific numbers of false negative and true positive calls, along with the proportion of the false negatives that consists of that TE family.

| TE | Total False Negatives | Candidate Region False Negatives | Proportion False Negatives | Total True positives |
|---|---|---|---|---|
| INE_1 | 11 | 10 | 0.12 | 0 |
| Gypsy_2_Dsim | 7 | 7 | 0.08 | 0 |
| F_element | 5 | 2 | 0.06 | 13 |
| FB4 | 5 | 5 | 0.06 | 15 |
| NOF | 5 | 4 | 0.06 | 0 |
| P_element | 5 | 4 | 0.06 | 28 |
| 1731 | 4 | 4 | 0.04 | 0 |
| Kepler | 4 | 4 | 0.04 | 0 |
| blood | 3 | 2 | 0.03 | 6 |
| BS | 3 | 2 | 0.03 | 3 |
| roo | 3 | 3 | 0.03 | 46 |
| 1360 | 2 | 2 | 0.02 | 0 |
| 297 | 2 | 1 | 0.02 | 15 |
| Copia | 2 | 2 | 0.02 | 14 |
| Copia2 | 2 | 2 | 0.02 | 0 |
| gypsy12 | 2 | 1 | 0.02 | 1 |
| HMS_Beagle | 2 | 0 | 0.02 | 2 |
| HMS_Beagle2 | 2 | 1 | 0.02 | 0 |
| 17_6 | 1 | 1 | 0.01 | 2 |
| accord | 1 | 1 | 0.01 | 8 |
| Bari1 | 1 | 0 | 0.01 | 1 |
| Blastopia | 1 | 1 | 0.01 | 8 |
| Circe | 1 | 1 | 0.01 | 0 |
| Doc | 1 | 0 | 0.01 | 8 |
| G4 | 1 | 1 | 0.01 | 0 |
| G6 | 1 | 0 | 0.01 | 1 |
| GATE | 1 | 1 | 0.01 | 0 |
| Gypsy_24_Dya | 1 | 0 | 0.01 | 4 |
| gypsy8 | 1 | 1 | 0.01 | 0 |
| Invader3 | 1 | 1 | 0.01 | 0 |
| Max_element | 1 | 0 | 0.01 | 2 |
| mdg1 | 1 | 0 | 0.01 | 9 |
| NewFam14 | 1 | 1 | 0.01 | 0 |
| pogo | 1 | 0 | 0.01 | 16 |
| Quasimodo | 1 | 0 | 0.01 | 3 |
| Rt1b | 1 | 1 | 0.01 | 4 |
| S_element | 1 | 0 | 0.01 | 1 |
| Stalker2 | 1 | 1 | 0.01 | 1 |
| Transpac | 1 | 0 | 0.01 | 6 |

| | | | | |
|---|---|---|---|---|
| 3S18 | 0 | 0 | 0 | 4 |
| 412 | 0 | 0 | 0 | 9 |
| BS2 | 0 | 0 | 0 | 5 |
| Burdock | 0 | 0 | 0 | 4 |
| diver | 0 | 0 | 0 | 1 |
| Doc6 | 0 | 0 | 0 | 5 |
| G_element | 0 | 0 | 0 | 1 |
| gypsy1 | 0 | 0 | 0 | 3 |
| gypsy6 | 0 | 0 | 0 | 1 |
| H | 0 | 0 | 0 | 25 |
| hopper | 0 | 0 | 0 | 1 |
| I_element | 0 | 0 | 0 | 8 |
| Idefix | 0 | 0 | 0 | 1 |
| Ivk | 0 | 0 | 0 | 5 |
| jockey | 0 | 0 | 0 | 44 |
| mdg3 | 0 | 0 | 0 | 3 |
| Nomad | 0 | 0 | 0 | 4 |
| Rt1a | 0 | 0 | 0 | 1 |
| Stalker4 | 0 | 0 | 0 | 1 |
| Tirant | 0 | 0 | 0 | 5 |

**Supplementary Table 2:** Numbers of true positive and false negative TEs, sorted by whether they are nested or not. Since short-read TE callers cannot distinguish multiple copies of the same TE family at one site, true positives and false negatives represent the number of nested TE groups rather than the total copy number.

| | Non-nested TE | TE nested with different family | Group of TEs nested with same family |
|---|---|---|---|
| True positives | 291 | 9 | 9 |
| False negatives | 54 | 28 | 8 |
| False negatives lost during candidate region detection | 39 | 20 | 8 |
